# Parvalbumin expression identifies subicular principal cells with high projection specificity

**DOI:** 10.1101/2024.08.13.606964

**Authors:** Gilda Baccini, Angelica Foggetti, Natalie Wernet, Karl-Alexander Engelhardt, Kerstin Kronenbitter, Jan Michels, Akos Kulik, Christian Wozny, Peer Wulff

## Abstract

The calcium-binding protein parvalbumin is an established marker for a subset of cortical inhibitory interneurons with similar biophysical features and connectivity. However, parvalbumin is also expressed in a small population of excitatory cells in layer 5 of the neocortex with specific sub-cortical projection targets. Parvalbumin may thus also in principal cells identify particular subclasses with distinct connectivity and function. Here we investigated whether parvalbumin is expressed in excitatory neurons of the hippocampal formation and if so, whether it delineated neurons with specific features. We report parvalbumin-expressing glutamatergic cells in the distal subiculum, which -based on location, connectivity and gene expression - separated into two subclasses: neurons in deep layers, which specifically project to the antero-ventral thalamus and neurons in superficial layers, which project to the mamillary bodies. Contrary to most adjacent pyramidal cells parvalbumin-positive neurons were non-bursting and displayed straight apical dendrites devoid of oblique dendrites. Functionally, the projections diverged from classical driver/modulator subdivisions. Parvalbumin expression thus marks two sub-types of subicular projection neurons with high target specificity and unique functional features.

## Introduction

The calcium-binding protein parvalbumin (PV) is one of the most well-known neuronal markers. PV itself modulates calcium-dependent metabolic and electric processes and shapes synaptic properties by modulating release kinetics and short-term plasticity^1–4^. As a neuronal marker, PV has been extensively used to identify a restricted subgroup of cortical inhibitory interneurons, characterized by particular electrophysiological properties as well as neuronal morphology and network connectivity^5–11^.

However, PV has also been found in a small population of excitatory pyramidal neurons in layer 5 of somatosensory, visual and motor cortex^12–18^. These neurons belong to a class of pyramidal cells called extra-telencephalic or pyramidal tract neurons, which have been reported to act as ‘drivers’ of feed-forward activity in subcortical areas^19,20^. Excitatory neurons expressing PV have long gone unnoticed with classical immunostaining possibly due to lower PV expression^17^ and currently, it is not yet clear whether, similar to GABAergic interneurons, PV-expressing excitatory cells may also constitute subpopulations of neurons with specific physiology, connectivity and function.

Recently, transcriptional profiling of neurons in mouse motor cortex has identified PV expression in specific populations of layer 5b. These PV+ neurons of taxonomic groups Npsr1 and Slc02a1^21,22^ displayed different levels of PV expression, distinct laminar distributions and projections as well as behaviour-related activity. Specifically, neurons containing high levels of PV transcript were located more superficially in layer 5b than neurons with lower PV content, projected to the thalamic premotor centre rather than the medulla, showed early and persistent rather than late preparatory activity, and appeared to control movement planning rather than initiation and termination^22^. PV expression may thus also identify subclasses of excitatory neurons with particular connectivity and function.

In this study we asked whether PV is also expressed in glutamatergic principal cells of the archicortical hippocampal formation and if so whether it would delineate subgroups of neurons with specific properties. PV expression has previously been reported by single cell transcriptomic analysis for a cluster of glutamatergic principal cells in the subiculum^23^. Such transcriptome-based clustering in the subiculum has been yielding regarding the identification of groups of neurons with common anatomical locations and projection areas^23–26^.

Here we show that PV-expressing principal cells are found in the medial to distal dorsal subiculum, where they occupy superficial or deep layers. These PV+ neurons project with high selectivity to either the mammillary body (MB) or the antero-ventral thalamus (AV), where they make excitatory connections with distinct synaptic properties. Analysis of available single cell transcriptome data confirmed the existence of two clusters of excitatory PV+ neurons, which separated into superficial and deep layers. These two clusters showed different levels of PV expression and displayed unique sets of expressed genes. In contrast to neighbouring principal cells in the distal subiculum, PV+ principal cells were non-bursting and showed unique morphological features. We thus propose that PV expression marks two new sub-types of subicular projection neurons with specific connectivity and functional features.

## Results

### A sub-population of parvalbumin-positive neurons in the distal subiculum is excitatory

PV is a prominent marker for a sub-group of inhibitory interneurons in the cortex^5,7^, but has been reported to also be expressed in a small subclass of glutamatergic pyramidal neurons in deep layer 5 of neocortex^16,17^. These layer 5 PV-positive pyramidal cells may comprise cell types with distinct gene expression, axonal projection and function^21,27^.

To explore whether PV+ glutamatergic neurons also exist in archicortical structures, we screened PV+ neurons in the dorsal hippocampus for co-expression of the vesicular glutamate transporters, VGLUT1 and VGLUT2, using fluorescent in situ hybridization (Fig. 1A). Whereas no co-localization of PV and VGLUT1 or 2 was apparent in dentate gyrus or the cornu ammonis, we did find a group of double labelled cells in the subiculum, where about 6 % of PV+ neurons co-expressed VGLUT2 and about 13% co-expressed VGLUT1 (Fig. 1A,B). Most of these PV+VGLUT+ cells were located in the deep (polymorphic) and superficial (pyramidal) layers of the distal subiculum (Fig. 1C). The expression of VGLUTs in PV+ neurons was surprising as in literature subicular PV+ neurons are considered GABAergic^28,29^. Indeed, double fluorescent in situ hybridizations confirmed that almost all PV+ cells co-expressed the vesicular GABA transporter (VGAT) (about 95%, 615 cells in 2 mice). Correspondingly we found a small percentage of VGAT+ cells to also express VGLUT1/2 (Supplementary Fig. 1A,B). Many of the VGLUT+VGAT+ cells were located in deep and pyramidal layers of distal subiculum, matching the location of the PV+VGLUT+ neurons (Supplementary Fig. 1C).

**Figure 1.**
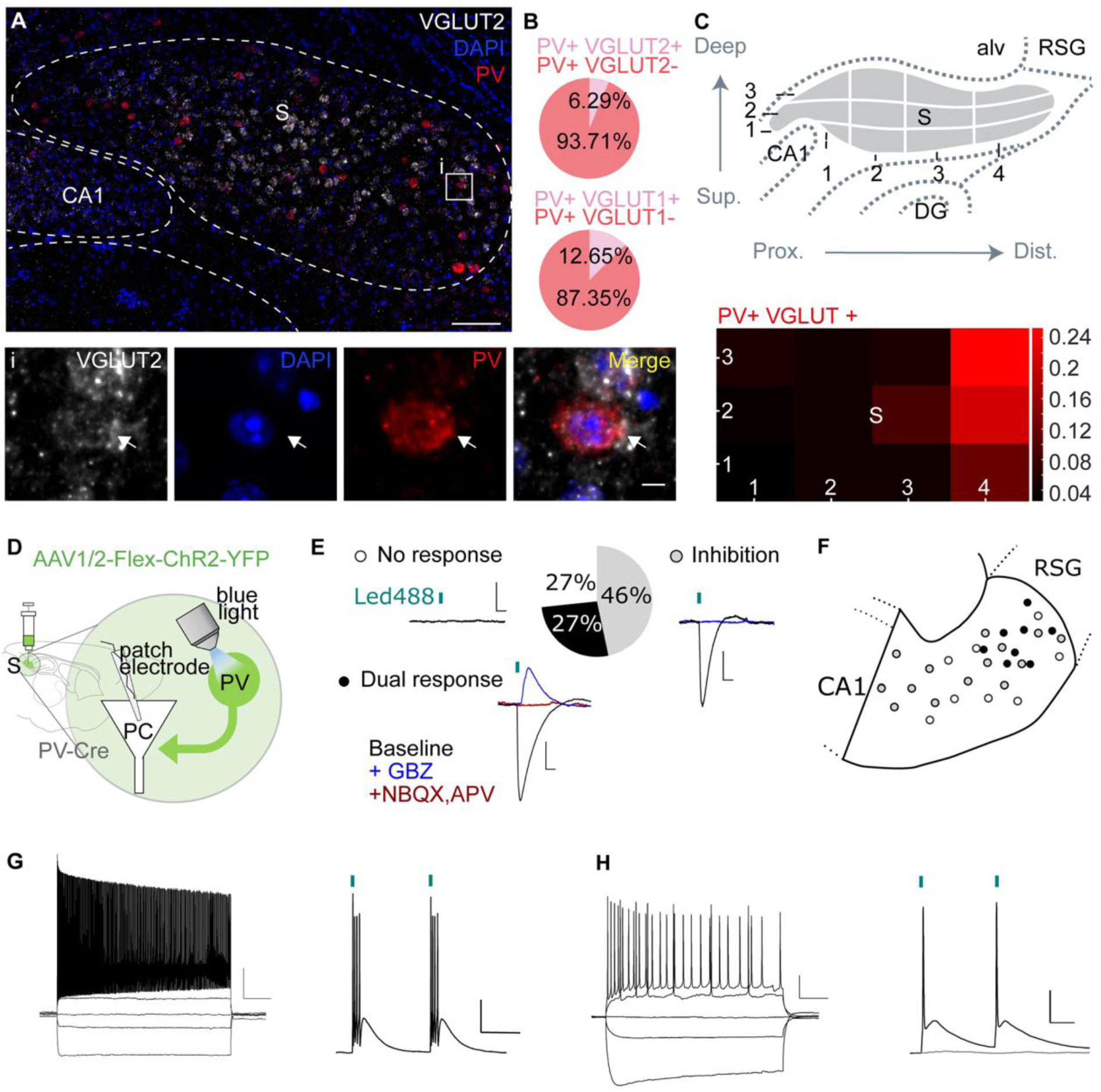
The distal subiculum contains parvalbumin-positive neurons that are excitatory. A) Example image of a double fluorescent in situ hybridization on a coronal brain section showing co-localization of VGLUT2 (white) and PV (red) mRNA in the subiculum. Nuclei are stained with DAPI (blue). Scale bar 100 µm. i, Magnification of the boxed area in A. The white arrow indicates a cell co-expressing PV and VGLUT2 mRNA. Scale bar 5 µm. B) A small percentage of subicular PV+ neurons co-expressed VGLUT2 (n=747 cells in 3 mice) or VGLUT1 (n=569 cells in 1 mouse) mRNA. C) To topographically describe the occurrence of VGLUT1/2 expressing PV+ neurons, we divided the long and short axes of the subiculum into 4 and 3 equal parts, respectively, to obtain 12 sub-fields along the proximal/distal axis and superficial/deep layers (upper panel). The relative occurrence of double labelled neurons is displayed in a heat map (double labelled cells per subfield/total double labelled cells, lower panel). CA1 cornus ammonis, DG dentate gyrus, S, subiculum; alv, alveus; RSG, retrosplenial cortex granular. D) Cartoon illustrating the electrophysiological recording of postsynaptic responses in pyramidal cells (PC, white) during optogenetic stimulation of PV+ neurons (green) in the subiculum of PV-Cre mice. E) 46% of recorded neurons showed exclusively inhibitory responses, 27% of cells showed both excitatory and inhibitory responses, 27% of recorded cells were unresponsive (n=30 cells in 5 mice). Example traces are displayed adjacent to the respective groups in the pie chart. Inhibitory responses could be blocked with gabazine (GBZ), excitatory responses could be blocked with NBQX and APV. Scale bar 2 mV, 50 ms. F) Cartoon depicting the location of the pyramidal cells receiving inhibitory input (gray), excitatory and inhibitory input (black) or no input (white) in the dorsal subiculum (cells are reported on a sagittal atlas plate 0.96 mm lateral (Paxinos and Franklin 2012). G, H) Current injections in CHR2-YFP-expressing PV+ neurons in distal subiculum revealed two different electrophysiological phenotypes: fast spiking, non-accommodating cells (G, left panel) and cells with slow and accommodating firing (H, left panel). Scale 20 mV, 200 ms. Both types of neurons showed high fidelity in their firing response following short pulses of light (right panels in G and H). Pulse duration 1 ms at 10% LED power; scale 10 mV, 50 ms at 10 Hz.

To probe whether VGLUT expression indeed indicated excitatory output, we analyzed post-synaptic responses after stimulation of PV+ neurons. To this end we expressed YFP-tagged channelrhodopsin-2 (ChR2-YFP) in subicular PV+ neurons by stereotaxic injection of Cre-dependent AAVs into the subiculum of PV-Cre transgenic mice. We then performed whole-cell patch-clamp recordings from non-labelled neurons along the proximo-distal axis of the subiculum (n= 30 cells in 5 mice) during light stimulation (Fig. 1D). While the majority of patched neurons (46%) exhibited only inhibitory responses, 27% of the neurons showed both excitatory and inhibitory postsynaptic currents (IPSC amplitude 152.3 ± 42.9, n=14, EPSC amplitude 13.0 ± 5.7 pA, n=8). Application of ionotropic receptor antagonists Gabazine (GBZ) or NBQX and APV, confirmed the GABAergic (n=15) and glutamatergic (n=5) nature of postsynaptic responses, respectively (Fig. 1E). Cells presenting with inhibitory and excitatory double responses had regular or bursting electrophysiological phenotypes (4 bursting, 1 regular, n=5) and were located towards the distal subiculum, where PV+VGLUT+ neurons had been detected (Fig. 1F). Targeted recordings of intrinsic properties from ChR2-YFP-expressing neurons in the distal subiculum (n=10) revealed two different electrophysiological phenotypes: 1.) fast spiking, non-accommodating cells, matching the properties of PV+ cortical interneurons (n=5; Fig. 1G)^30,31^, 2.) cells that displayed a slow and accommodating action potential firing, like it has been described for pyramidal neurons (n=5; Fig. 1H)^32–35^.

We thus identified a sub-population of PV+ neurons in the distal subiculum, which express vesicular glutamate transporters, provide excitatory out-put to neighbouring principal cells and likely show electro-physiological properties reminiscent of pyramidal neurons.

### Subicular PV-positive principal cells project selectively to the anterior thalamus and the mammillary body

The subiculum is the main output station of the hippocampal formation and its principal cells send projections to several cortical and subcortical targets^26,36^. To test whether the PV+ excitatory neurons we identified also project outside the subiculum, we fluorescently labelled PV+ subicular neurons using Cre-dependent AAVs in PV-Cre transgenic mice. Indeed, many labeled axons were leaving the subiculum through the corpus callosum and joined the post-commissural fornix to reach thalamic and hypothalamic regions. Interestingly, no fibers were detected in the internal capsule or the reticular thalamus (RT) which have been described as parts of a parallel collateral path for subicular projections to subcortical target areas^37–39^ (Fig. 2A).

**Figure 2.**
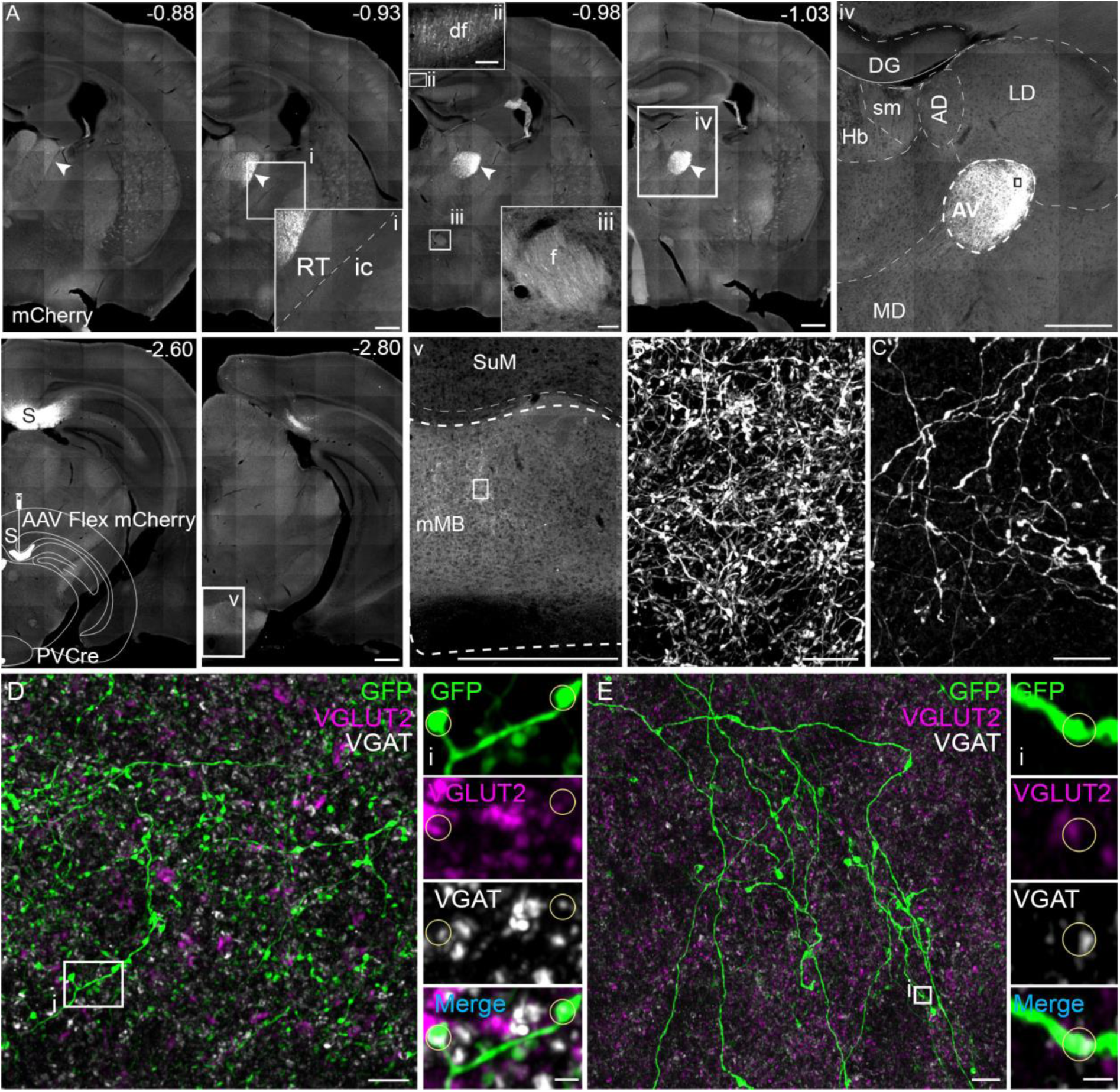
Glutamatergic PV+ cells send projections to the anterior thalamus and the mammillary body. A) Coronal brain sections in rostro-caudal sequence depicting the pathway of PV+ axonal projections in a PV-Cre mouse injected with mCherry-expressing Cre-dependent rAAVs (AAV-FLEX-mCherry) in the dorsal subiculum. Insets i, ii and iii show magnifications of the boxed areas demonstrating the absence of fibers from the internal capsule (ic) and the reticular thalamus (RT) but their presence in the dorsal fornix (df) and fornix (f), respectively. iv, close up of the boxed area in the preceding image (bregma level -1.03) showing the high density of labeled fibers in the antero-ventral thalamus (AV). v, magnification of the boxed area in the preceding image (bregma -2.80), showing some mCherry-positive fibers also in the medial mammillary nucleus (mMB). Scale bar 500 µm in all overview images, i -iii scale bar 50 µm, iv and v scale bar 500 µm. B,C) Close up of the boxed areas in iv,v, respectively, showing the dense network of mCherry-positive fibers in the AV and sparse positive fibers in the mMB. Scale bar 10 µm. Note the injection site in the Subiculum (S) is visible at bregma level -2.60. D,E) Images from the AV (D) and the mMB (E) of a PV-Cre mouse injected with a GFP-expressing Cre-dependent rAAV in the dorsal subiculum, stained for GFP (green) as well as the presynaptic markers VGLUT2 (purple) and VGAT (white). Images on the right show magnifications of the boxed areas on the left. Circles indicate boutons immunoreactive for GFP, VGLUT2 and VGAT. Image stacks are 2.5 µm in D and 0.6 µm in E. Scale bar 5µm in the overview, 1 µm in the magnifications. AD, antero-dorsal nucleus of the thalamus; AV, antero-ventral nucleus of the thalamus; DG, dentate gyrus; df, dorsal fornix; f, fornix; Hb, habenula; ic, internal capsule; LD, lateral dorsal nucleus of the thalamus; MD, mediodorsal nucleus of the thalamus; mMB, medial mammillary nucleus; RT, reticular thalamus; sm, stria medullaris of the thalamus; S, Subiculum; SuM, Supramammillary nucleus.

Axonal terminals were highly confined to the antero-ventral thalamus (AV) and the mammillary body (MB) (Fig 2). In the AV, we found dense nets of fibers along the entire rostro-caudal axis, which were particularly pronounced in the lateral part (Fig. 2A,B, Supplementary Fig. 2). No axonal projections were detected in other parts of the anterior thalamus. Within the MB labelled fibers were less dense and restricted to the medial part (mMB) (Fig. 2A,C). No obvious labeling was found in other known target regions of the subiculum (Supplementary Fig. 2)^26,36^. To confirm that the PV+ projections we observed were not contaminated by accidental transduction of neighbouring pre-subicular neurons, we selectively transduced PV+ neurons in the pre-subiculum (Supplementary Fig. 3). However, axonal arborizations were locally restricted and no GFP+ axon terminals were evident in AV or MB. To confirm the peculiarity of the projection tract and the target selectivity of PV+ subicular projection neurons, we stereotactically injected AAVs expressing GFP under the control of the CaMKII promoter (AAV-CaMKII-GFP) to non-specifically label principal cells in the subiculum. As expected, tracing of GFP+ axons showed fiber projections branching into both fornix and internal capsule forming dense nets of terminals throughout all parts of the anterior thalamus and the MB (Supplementary Fig. 4) as well as in the RT and other known target regions^26,36,40^. To confirm synaptic contacts of PV+ subicular projections in AV and mMB and to determine their neurochemical nature we immunostained for excitatory and inhibitory presynaptic markers. GFP+ fibers in AV and mMB showed a similar occurrence of VGAT-and VGLUT2-positive boutons and both markers frequently co-localized (about 93% and 59% of VGLUT2+ puncta co-localized with VGAT in AV and mMB, respectively Fig. 2D,E).

In summary our tracing experiments revealed that PV+ neurons in the subiculum project exclusively through the fornix to target the lateral AV and to a lesser extent the medial MB. In agreement with the in situ data we find that these neurons express vesicular transporters for both GABA and glutamate.

### PV+ subicular projections form excitatory synapses in AV and MB

To investigate whether PV+ subicular projections formed functional synapses and whether these were excitatory or inhibitory we expressed ChR2-YFP in PV+ neurons of the subiculum and made voltage-clamp recordings from cells in the AV in acute sagittal or coronal slices (Fig 3, Supplementary Fig. 5). Light-evoked responses were recorded at different potentials to separately monitor EPSCs and IPSCs, respectively (Fig. 3B). Out of 160 neurons from sagittal and 57 neurons from coronal slices, 43 and 19 cells, respectively, showed excitatory responses. However, none of the cells showed inhibitory responses, neither direct, nor di-synaptic (feed forward inhibition) (Fig. 3, Supplementary Fig. 5). Current clamp recordings showed firing properties typical for thalamic neurons, irrespective of whether they responded to optogenetic stimulation or not (n=22)^41^ (Supplementary Table 1). Recorded cells were filled with neurobiotin and mapped onto the 3D volume of the AV (Fig. 3C, Supplementary Fig. 5). Light responsive neurons were located in a confined volume in the lateral zone of the dorso-caudal AV, corresponding to the terminal field of YFP+ fibers (Fig. 3C, Supplementary Fig. 5). Synaptic currents reached half-maximal values at 12.5% and maximal values at 30% (n=9) (Fig. 3D). Stimulation at 30% showed time-locked postsynaptic responses with fixed and short latency (3.6 ± 0.1 ms; n=38) and ranged from 2 to 77 pA in amplitude (Fig. 3E,F)). We confirmed the dependency on presynaptic action potentials by application of TTX (tetrodotoxin), which abolished postsynaptic responses (n=6). Addition of the potassium channel blocker 4-AP (4 aminopyridine) rescued the light-evoked EPSCs, corroborating the monosynaptic nature of the synaptic response (n=6). EPSCs were mediated by ionotropic glutamate receptors as they were abolished by blockers of AMPA and NMDA receptors, NBQX and APV, respectively (n=6) (Fig. 3G).

**Figure 3.**
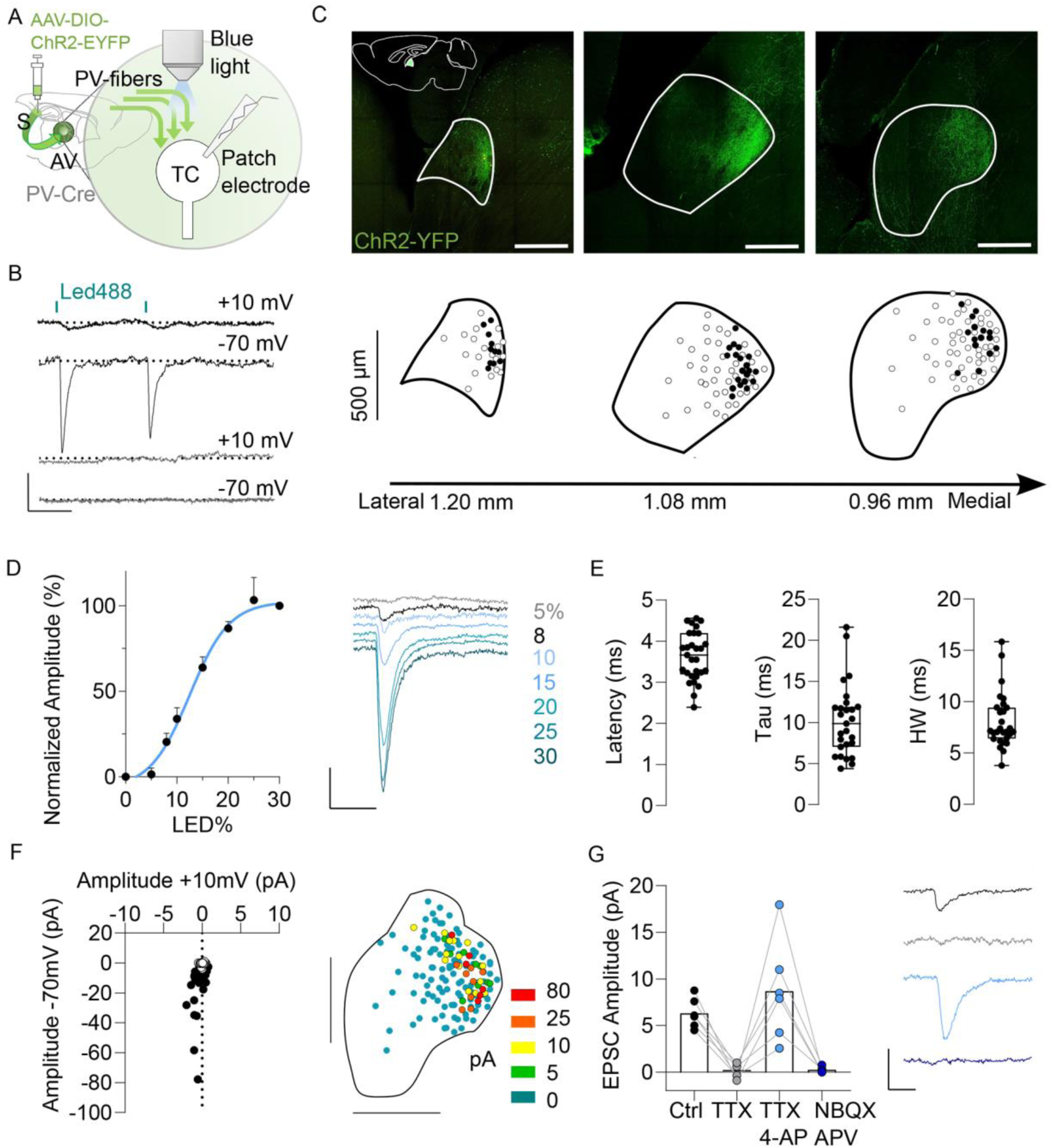
Subicular PV-positive principal cells form functional excitatory synapses onto AV neurons. A) Cartoon illustrating the experimental setup to record postsynaptic responses in thalamic cells (TC) of the AV during photostimulation of PV+ ChR2-YFP-expressing fibres from the subiculum (green). B) Example traces of current responses for a responsive (upper traces) and a non-responsive (lower traces) TC held successively at -70 and +10 mV to record EPSC and IPSC, respectively. Response latencies were measured from the onset of the light pulse (blue bars, 470 nm, 1 ms, 20% LED power corresponding to 4 mW/mm^2^). Scale bar 10 pA, 50 ms. C) Top, confocal image of a sagittal slice showing the location of the ChR2-YFP positive fibres (green) in the dorso-caudal AV. Bottom, whole-cell recorded neurons were filled with neurobiotin and registered onto sagittal planes (black dots, EPSC; white dots, no response). D) Left, mean input–output curve of EPSC amplitude as a function of light intensity (n=9). Results are means ± s.e.m. Right, example of current responses of a TC to optogenetic stimulation of PV+ afferents at different intensities. Scale bar 5 pA, 20 ms. E) Box-and-whisker plots showing latency, weighted decay time constant and half-widths of EPSCs (LED 30% corresponding to 5.6 mW/mm^2^, n=28). F) Left, response amplitudes measured at -70 mV and +10 mV. Right, map illustrating the location of all patched cells in the AV. EPSC response amplitudes are colour-coded. I) Pharmacological analysis of recorded EPSCs in the AV with example traces depicted in the corresponding color shown on the right. Light stimulated responses are blocked by TTX (grey), but are recovered by 4-AP (light blue). Responses are suppressed by NBQX (25 µM) and APV (100 µM) (dark blue). Scale bar 10 pA, 25ms.

Photo stimulation of PV+ ChR2-expressing fibres in the mMB evoked EPSC in 7% (3 out of 44) of patched cells (amplitude: 19.5 ± 13.5 pA, latency: 3.5 ± 3 ms, n=3), but again, none responded with inhibitory responses (Supplementary Fig. 6). Application of TTX and 4-AP (n=2) confirmed that EPSCs also in the mMB were action potential-dependent and monosynaptic (Supplementary Fig. 6).

These data demonstrate the formation of functional glutamatergic synapses in AV and MB. However, we did not observe any inhibitory responses, despite the expression of VGAT in PV+ projection neurons. Indeed, also in the presence of the glutamate receptor antagonists APV and NBQX no inhibitory responses could be observed in the AV, neither by single light-stimuli (1 ms, 30%, n=11 cells with and n=14 cells without EPSCs) nor by trains of stimuli at different frequencies (20 Hz, n=4 with and 4 without EPSCs, 40 Hz and 100 Hz, n=5 with and 5 without EPSCs per protocol, Supplementary Fig. 7). We also found no indication of light-evoked responses mediated by extra-synaptic GABA-A receptors or GABA-B receptors (Supplementary Fig 7). To further probe the potential inhibitory phenotype of PV+ projections we injected a distal-less homeobox 5 and 6 (Dlx) enhancer-driven rAAV into the subiculum of PV-Cre:Ai9 transgenic mice to express ChR2-YFP in GABAergic PV+ subicular neurons^42,43^. In agreement with our electrophysiological results no labeling was detected in AV or MB (Supplementary Fig. 8).

From these experiments we conclude that PV+ projections make monosynaptic connections to spatially confined neuronal groups in AV and to a lesser extent in MB, which are both excitatory in nature. In contrast to known efferents from parahippocampal regions these projections do not recruit feed-forward inhibition via the reticular thalamus^38,44^.

### PV+ subicular projections to AV display driver-like features

Cortico-thalamic projections have been classified into “drivers” and “modulators” for the sensory thalamus^20^. Here drivers target proximal dendrites with large synaptic boutons, show high release probability and elicit large amplitude, ionotropic responses, which change “all or none” with increasing stimulation^45^. Drivers reliably evoke action potentials and thus faithfully transmit information. In contrast, modulators target distal dendrites with small synapses, exhibit low release probability and evoke smaller ionotropic responses, which change gradually in amplitude with increasing stimulation. Enhanced stimulation additionally recruits metabotropic responses. Modulators fine-tune the information relayed by thalamic cells (reviewed in^46,47^). Basic features distinguishing drivers and modulators are present in all thalamic nuclei but may vary in expression^48^.

To assess distinguishing features for PV+ subicular afferents to the AV, we first studied release properties in a paired-pulse protocol. We found that the paired pulse ratios (PPRs) remained below 1 at all interstimulus intervals tested (50 to 300 ms), indicating synaptic depression (Fig. 4A,B). This was independent of the light intensity as tested during 10 Hz stimulation (Fig. 4C). Such paired pulse depression suggests high release probability, which is deemed a driver feature. However, unlike a classical driver, PV+ inputs showed a graded response with gradually increasing EPSPs upon enhanced stimulation^45,49^ (Fig. 3D). Driver projections are able to evoke action potentials (APs) in postsynaptic thalamic neurons^50^. To test whether PV+ projections would in principle be able to do so, we depolarized thalamic neurons to -50 mV, where they tend to fire with tonic discharges, and stimulated subicular PV+ afferents with light. We found that 3 out of 5 tested cells responded with AP generation (Fig. 3D). To determine whether PV+ projections would recruit postsynaptic metabotropic glutamate receptors of group I (mGluRI) we stimulated PV+ projections at 100 Hz for 200 ms^45,51^, while recording from thalamic neurons which showed ionotropic responses (n=6). However, we found no indication of mGluRI-mediated responses (Fig. 4E).

**Figure 4.**
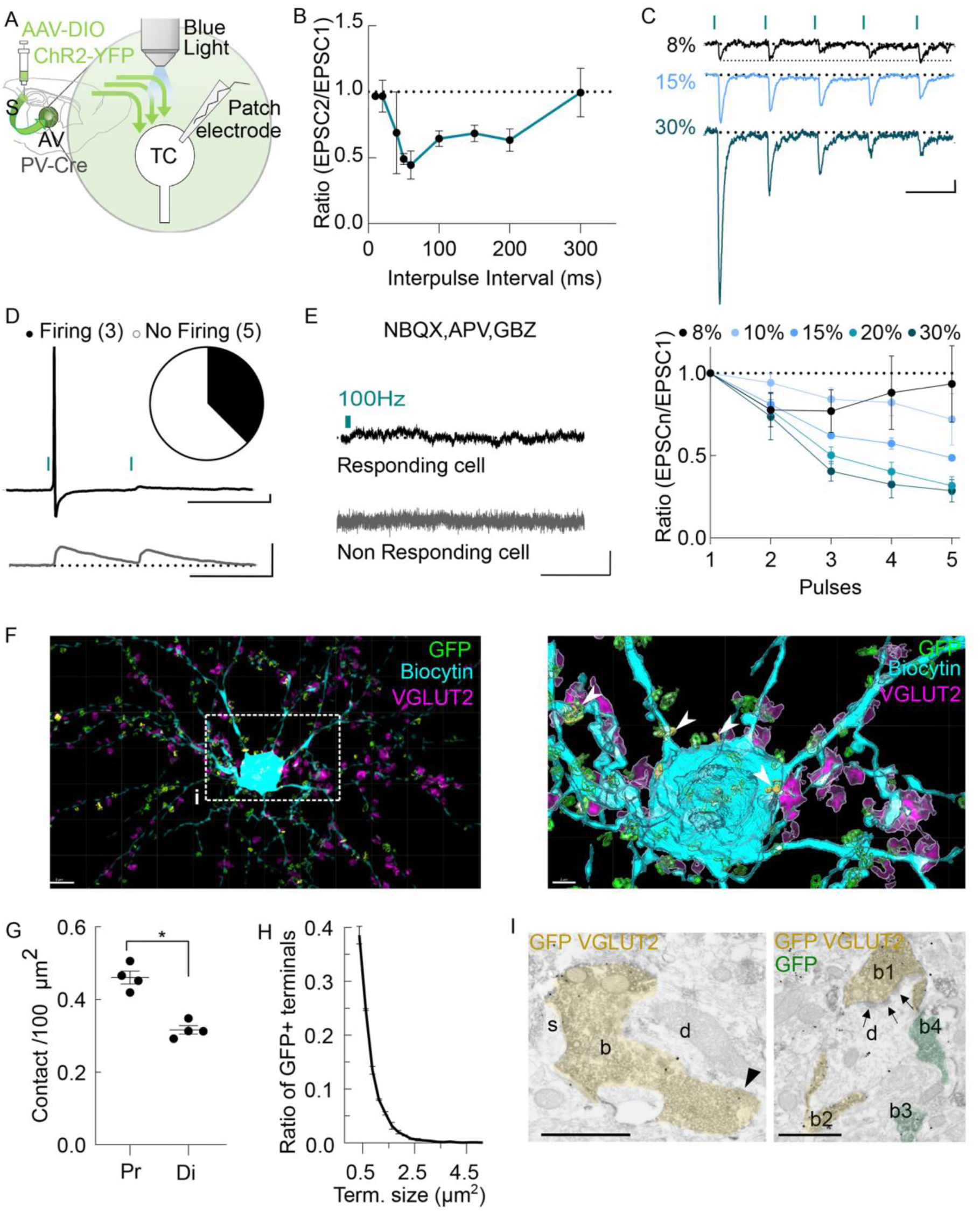
Subicular PV-positive principal cells establish connection in AV, that diverge from classical driver/modulator categories. A) Illustration of the electrophysiological recording set up. B) Paired pulse ratios (PPRs) of EPSCs at different interstimulus intervals indicated that PV+ projections in the AV display synaptic depression (n= at least 4, for 100 ms n=18, 150 ms n=10, 200 ms n=8). Data are shown as mean ± sem. C) A train of stimuli at 10Hz shows synaptic depression at different light intensities of stimulation (minimal stimulation at 8% corresponds to 1.7, maximal at 30 to 5.6 mW/mm^2^). The top shows example traces, the bottom EPSP ratios. Scale bar 5 pA, 100 ms. D) Example traces of (top) a cell that responds with action potential firing and (bottom) a cell that does not show an action potential response. When held at -50mV and stimulated with light, 3 out of 8 cells responded with action potential firing. Scale bar 5 mV, 100 ms. E) Example voltage clamp recordings in a cell that showed light-evoked EPSCs (top, n=5) or did not (bottom, n=3) during high frequency stimulation (100 Hz, 1 ms pulse duration, 30% LED power, held at -50 mV) in the presence of ionotropic glutamate and GABA antagonists (NBQX, APV and GBZ) showed no indication of metabotropic responses. Scale bar 5 pA, 1 s. F) Left, 3D isometric view of a thalamic neuron filled with biocytin (blue) and presynaptic VGLUT2-(purple) and GFP-(green) immunoreactive buttons of subicular projections. Right, 3D-surface reconstruction of the neuron on the left at higher magnification. Only VGLUT2-and GFP-positive surfaces with a distance of 0.5 µm from the filled neuron are depicted. Colocalization of VGLUT2 and GFP is shown in yellow (arrowheads). G) Putative synapses were found more frequently on proximal dendrites and somata (Pr) than on distal dendrites (Di). p=0.0111, paired t-test. H) The size of GFP+ presynaptic terminals showed considerable variation. I) Electron micrographs showing synaptic contacts (arrow head and arrows) between GFP-positive (peroxidase reaction product) and VGLUT2-positive (immunoparticles) boutons (b, b1, b2) and large calibre dendritic shafts (d) of thalamic neurons. S, dendritic spine; b3 and b4, GFP-positive but VGLUT2-negative axon terminals. Scale bar 1 µm.

Finally, to also assess morphological features of PV+ subicular afferents, we examined the size and distribution of presynaptic terminals. Putative boutons were identified by colocalization of YFP and VGLUT2 immunoreactivity in close proximity (0.5 µm) to light-responsive reconstructed thalamic neurons (n=4) using high-resolution confocal microscopy. We found the density of putative presynaptic terminals to be higher on proximal dendrites and somata than on distal dendrites (Fig. 4 F,G). Sizes of presynaptic terminals ranged from 0.3 to 4.76 µm^2^ (mean: 0.80±0.07 µm^2^) (Fig. 4H). Immunoelectron microscopic analysis showed GFP+ terminals predominantly targeting dendritic shafts and occasionally dendritic spines (which were frequently immunoreactive for VGLUT2 (Fig. 4I) and in randomly selected sections had a maximum cross section area of 2 µm^2^ (Fig. 4I, Supplementary Fig. 12).

In summary, PV+ subicular projections to the AV formally meet several criteria of driver type neurons but show smaller EPSC amplitudes, which change gradually with stimulation intensity, suggesting that they diverge from the classical driver/modulator classification.

### PV+ excitatory neurons in the distal subiculum are regular spiking and project either to AV or MB

Projecting neurons in the subiculum are organized topographically along the proximo-distal and dorso-ventral axis^23,36,52^. In addition, subicular neurons are divided into two main classes based on their firing properties: regular firing neurons, located towards the proximal and burst firing neurons, located towards the distal subiculum^35,53,54^.

To determine the anatomical distribution of PV+ subicular projection neurons, we injected the retrograde tracer fast blue (FB), into either AV (n=3 mice) or MB (n=4 mice). FB+ neurons projecting to AV were located in deep layers of the distal subiculum whereas FB+ neurons projecting to MB were located in more superficial layers of the medial to distal subiculum^55,56^ (Fig. 5A,B). Similarly, FB+ neurons immuno-reactive for PV projecting to AV were found mainly in deep layers, PV+ neurons projecting to MB were located in more superficial layers of the distal subiculum (Fig. 5A,B; Supplementary Fig. 9) suggesting that PV+ principal cells constitute two adjacent populations with laminar separation. PV+ neurons retrogradely traced from the AV expressed significantly less PV and had smaller pyramidal cell-like somata when compared to neighbouring FB-PV+ putative interneurons (Fig. 5C,D).

**Figure 5.**
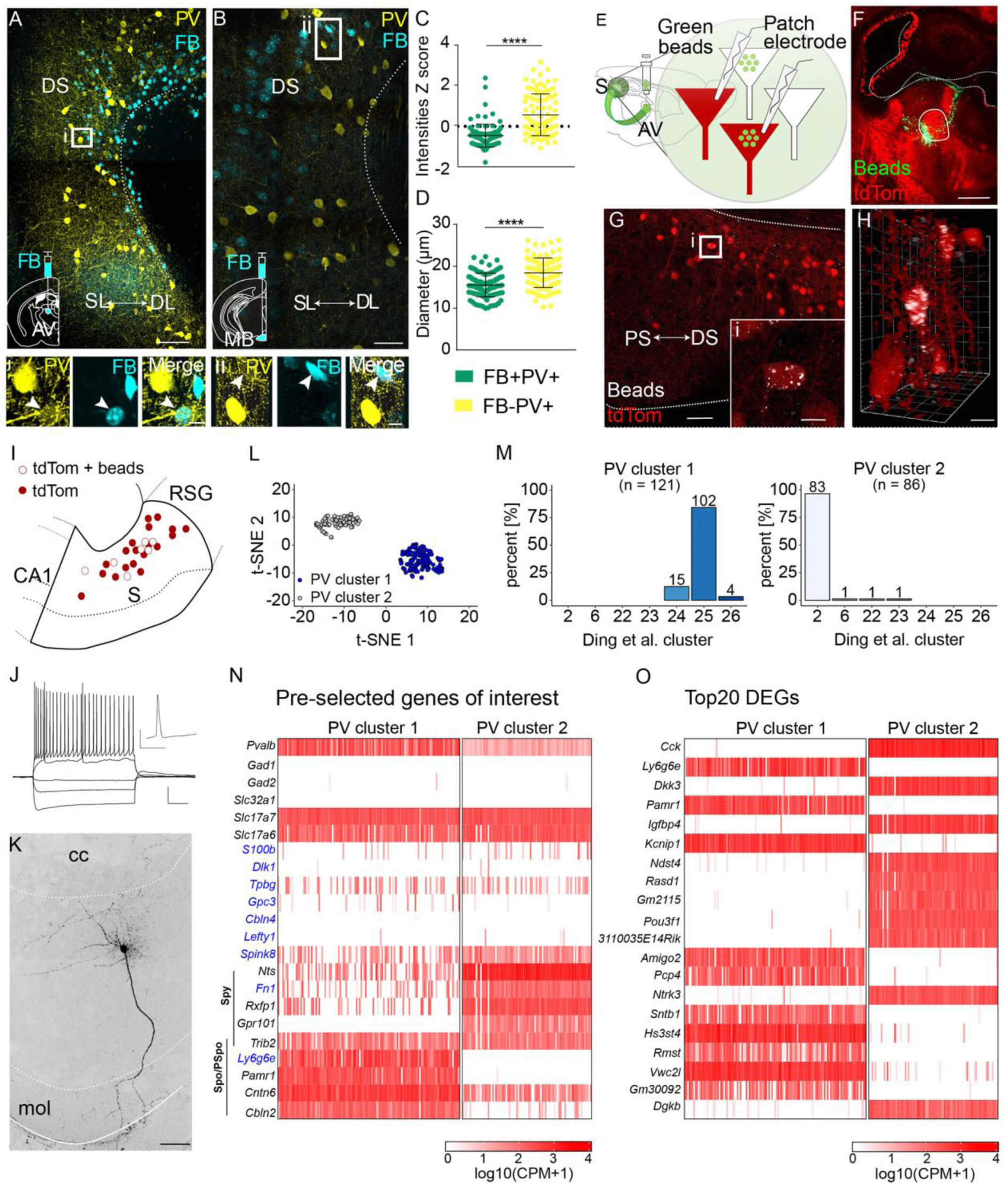
PV+ principal cells of the distal subiculum belong to two distinct transcriptomic clusters, target either AV or MB and are regularly firing. A) Confocal image of the subiculum of a wild-type mouse injected with fast blue (FB) in the AV, showing retrogradely labelled cells (blue) immuno-positive for PV (yellow). Scale bar 100 µm. i, Higher magnification of boxed area in A, showing one neuron which is both PV and FB positive (white arrowhead). Scale bar 10 µm. B) Image of the distal subiculum of a wild type-mouse injected with FB in the MB, displaying retrogradely labelled cells immunoreactive for PV (yellow). Scale bar 50 µm. ii, Higher magnification of boxed area in B, showing colocalization of PV and FB in one neuron (white arrowhead). Scale bar 10 µm. C) PV expression in PV+ cells that were retrogradely traced by FB is lower than in FB-negative PV+ cells. Each dot represents one cell, normalized by ZScore (lme p=9.28-16). D) FB+ PV+ neurons are smaller in diameter than FB-PV+ cells (lme p= 1.13e-9). E) Recording configuration, tdTomato-positive neurons were recorded in the subiculum seven days after injection of green retrobeads into the AV of PV-Cre::Ai9 mice. F) Brain slice showing the injection site of green retrobeads into the AV of a PV-Cre::Ai9 mouse. G) Image of the subiculum in the same mouse. Scale bar 50 µm. i, Higher magnification of the boxed cell in G, showing colocalization of tdTomato and retrobeads. Scale bar 10 µm. H) 3D perspective of some cells in G, with two tdTomato+ cells containing beads. Scale bar 5 µm. I) Cartoon illustrating the distribution of recorded tdTomato+ neurons with pyramidal cell like morphology, some of which contained beads. Patched cells were located in the deep layers and were more numerous toward the distal subiculum. J) Left, example of a recording from a tdTomato+ neuron containing beads showing voltage responses to increasing current injections (n=8). Scale bar 20 mV, 20 0ms. The inset shows the first action potential of the trace at rheobase +50 pA at higher temporal resolution. Scale 40 mV, 10 ms. K) Image of the neuron recorded in J filled with biocytin. Note the large apical dendrite devoid of oblique dendrites. Scale bar 100 µm. L) tSNE plot of the two clusters of PV-positive cells (filled circles PV cluster 1, open circles cluster 2). M) 102 cells of PV cluster 1 (in total n=121 cells) belong to group 25 identified by Ding et al.^23^, and 83 cells of PV cluster 2 (in total n=86 cells) belong to group 2. N) Selection of genes that are differentially expressed in cluster 1 and 2 including marker genes identified by Cembroswki et al., 2018^24^, shown in blue. O) Top 20 differentially expressed genes (DEGs) between cluster 1 and cluster 2. CA, cornu ammonis; cc, corpus callosum, DL, deep layer; DS, distal subiculum; mol, molecular layer, PS proximal subiculum, RSG, retrosplenial cortex granular; S, subiculum; SL, superficial layer.

To probe the firing properties of PV+ principal cells, we focused on the population projecting to the AV. We injected green fluorescent retrobeads into the AV of PV-Cre::Ai9mice, in which all PV+ neurons express tdTomato. This allowed us to identify PV+ projection neurons by co-localization of tdTomato and beads for targeted whole cell recordings (Fig. 5E-I). Current injection above threshold or at 50 pA above the rheobase^35,57^ never elicited bursting and the initial firing frequency always remained below 100 Hz (Fig. 5J; Supplementary Fig. 10; n=8), identifying these neurons as regular firing^53,57,58^. Recordings from tdTomato+ cells with pyramidal soma morphology in deep and more superficial layers of distal subiculum of animals not injected with beads showed the same regular firing phenotype (n=18; Supplementary Fig. 10). These recordings may have included PV+ principal cells that project to the MB. In contrast neighbouring tdTomato-negative pyramidal neurons showed typical burst firing (n=5; Supplementary Fig. 10). Analysis of additional sub-and supra-threshold properties also distinguished tdTomato+ from neighbouring tdTomato-negative pyramidal cells (Supplementary Fig. 10). Biocytin filling of recorded tdTomato+ neurons revealed a pyramidal cell shaped soma with basal dendrites emerging from the soma and a straight larger apical dendrite extending into superficial layers and ending in a tuft (n=10). Unlike typical regular or burst-firing pyramidal cells of the subiculum the apical dendrite was almost devoid of oblique dendrites (Fig. 5K; Supplementary Fig. 10;^57^).

PV+ subicular projection neurons to AV and MB thus lie in deep and more superficial layers of distal subiculum, respectively. Unlike the majority of distal subicular neurons, PV+ principal cells were regular firing and displayed straight unbranching apical dendrites that ended in a tuft in the superficial layer. We thus suggest that these neurons form a specific type of subicular pyramidal cell.

### PV+ glutamatergic neurons can be grouped into two transcriptomically distinct clusters within distal subiculum

We identified a population of PV+ excitatory neurons in the distal subiculum, which could be divided based on anatomical location and projection targets: a main subgroup residing in deep layers projecting to the AV and a smaller subgroup located in more superficial layers projecting to the MB. Consistently, Ding et al. (2020)^23^ recently identified several transcriptomically distinct glutamatergic cell types with defined anatomical locations using single-cell RNAseq in the subiculum. We re-analyzed these data to determine how PV+ excitatory neurons mapped onto the transcriptomically defined cell types, focusing on the SMARTseq samples provided by Ding et al. (2020), containing 1081 sequenced glutamatergic neurons. We found that PV expression was largely restricted to samples belonging to two clusters, cluster 2 (moderate PV expression) and 25 (strong PV expression)^23^ both located in the distal subiculum (Supplementary Fig. 11). Whereas cluster 2 expressed marker genes for the superficial (pyramidal) layer (e.g. *Nts*, *Fn1* or *Gpr101*), cluster 25 expressed markers for the deep (polymorphic) layer (e.g. *Ly6g6e* or *Pamr1*)^23^, matching the anatomical location and laminar distribution of the MB and AV projecting cells identified in this study.

We next selected all PV+ neurons from the SMARTseq samples (*Pvalb* CPM threshold ≥ 5, resulting in 207 PV+ cells), re-clustered them, and determined how they map onto the previously identified cell types. Identification of variable genes, dimensionality reduction, and clustering was performed in R using the Seurat package. This re-clustering resulted in two groups of PV+ neurons that closely corresponded to the clusters 2 and 25^23^ (Fig. 5 L,M). PV cluster 1 comprised 121 cells and mostly overlapped with cluster 25 (84.3% overlap). By contrast, PV cluster 2 comprised 86 and almost exclusively overlapped with cluster 2 (96.5% overlap). PV cluster 1 displayed strong PV expression and abundantly expressed markers of superficial layers, whereas PV cluster 2 showed moderate PV expression and abundantly expressed markers of deeper layers (Fig. 5N). These results support the idea of two subgroups of PV+ glutamatergic projection neurons residing in neighbouring but distinct layers of the distal subiculum.

To test whether these two PV clusters can be distinguished further by their transcriptome profiles, we performed a differential gene expression analysis and extracted the top 20 differentially expressed genes (DEGs) based on mean log2 fold change values. We found that PV clusters 1 and 2 express unique sets of DEGs with *Kcnp1*, *Amigo2*, *Pcp4*, *Sntb1*, *Hs3st4*, *Rmst*, *Vwc2l*, and *Gm30092* strongly expressed in PV cluster 1 and *Cck*, *Dkk3*, *Igfbp4*, *Ndst4*, *Rasd1*, *Gm2115*, *Pou3f1*, *3110035E14Rik*, *Ntrk3*, and *Dgkb* strongly expressed in PV cluster 2 (Fig. 5O; see discussion).

Accordingly, the re-analysis of available transcriptomic data from the subiculum supports our finding of two separate populations of PV-expressing pyramidal cells with adjacent but distinct anatomical locations and show that these 2 types of target-specific projection neurons express unique sets of genes.

## Discussion

We have identified excitatory PV+ neurons in the distal subiculum with morphological and electrophysiological properties of pyramidal cells. Based on location, connectivity and gene expression data these PV+ excitatory neurons can be divided in to two subclasses: neurons with strong PV expression in deep layers, which send strong and specific projections to the antero-ventral thalamus and neurons with moderate PV expression in superficial layers, which send sparser projections to the medial mamillary bodies. Focusing on the main group of AV targeting neurons we found that PV+ neurons projected exclusively through the fornix making monosynaptic connections with spatially confined neuronal groups without recruiting feed-forward inhibition via the reticular thalamus. Functionally, these projections diverge from classical driver/modulator subdivision. Unlike neighbouring pyramidal cells in the distal subiculum PV+ neurons showed a non-bursting phenotype and displayed straight and unbranching apical dendrites.

In cortical areas, parvalbumin expression is a prominent marker for a small group of GABAergic interneurons comprising basket cells, chandelier cells and bistratified cells with homogeneous biophysical features and specific circuit connectivity^5,7,59^.

However, occasional reports have indicated that PV is also expressed in a small population of excitatory cells of the neocortex in adult rodents and primates, although at lower levels^17,18,60^. This finding has recently gained support from single cell transcriptome analysis of neocortical neurons^21^. In cortical pyramidal cells PV expression seems to be restricted to extra-telencephalic neurons (ET) of layer V (also known as pyramidal tract neurons) in somatosensory, motor, auditory and visual cortex and to a smaller group of neurons in layer 6b^18,21^.

We tested for PV expression in glutamatergic cells of the hippocampal formation and report PV+ pyramidal neurons in the distal part of dorsal subiculum. Based on projection targets and laminar distribution these PV+ neurons separated into two sub-groups. PV+ neurons located in the deep layers that selectively projected to the AV and PV+ neurons located in more superficial layers projecting exclusively to mMB. Laminar location and projection specificity are consistent with previous studies, which reported that subicular outputs to thalamus and mamillary bodies are of different laminar origin^55,56,61^. Specifically, projection neurons targeting AV and AM were located in deeper layers of distal subiculum^55,62^, projection neurons targeting the mamillary body were described in the superficial central part of the subiculum comprising subpopulations with or without collaterals to retrosplenial cortex or entorhinal cortex^26,55,63^. A more general classification suggested that dorsal subiculum neurons fall into four groups: 1.) Distal neurons with local collaterals and projections to hypothalamus, retrosplenial cortex or both areas; 2.) Cells located more centrally, which send their axons to the anterior thalamic nuclei; 3.) Proximal cells projecting broadly to multiple targets, including the nucleus accumbens, lateral septal nucleus, thalamus, periaqueductal grey, and hypothalamus^24,25,64^; 4.) Neurons, distributed across the dorsal subiculum with local collaterals and rare long-range axons. The PV+ pyramidal neurons described here likely form two subgroups within this system of subicular projection neurons with collaterals in the subiculum and very specific projection targets in AV and MB. Re-analysis of available subicular single cell transcriptome data^23^ supports this notion. We identified two populations of PV+ excitatory neurons. PV cluster 1, showed higher *Pvalb* transcript levels and expressed *Ly6g6e*, a marker gene for deep layers of subiculum, likely identical with those PV+ neurons that project to AV^26^. PV cluster 2 with lower levels of *Pvalb* transcripts, which abundantly expressed *Fn1*, a marker gene for the pyramidal layer of subiculum (SUB-Fn1), likely identical with the PV+ neurons projecting to MB of the superficial layer.

The transcriptional profile assigns PV cluster 1 to cluster SUB-NP (*Ly6g6e*, *Cntn6*, *Pamr1*, *Cbln2*, and *St3gal1*) described by Ding et al., 2020^23^, which show regional similarity with layer 5/6 neurons in isocortex. PV+ neurons of cluster 2 on the other hand belong to the SUB-ProS cluster (*Fn1*, *Pou3f1*, *Bcl6*, *Bhlhe40*, *Blhlhe22*), which are similar to layer 5 cells in isocortex. Interestingly, none of our two PV clusters seems part of the SUB-CT cluster (Rorb), comprising corticothalamic cells related to L6 CT (*Syt6*, *Rasgrf2*, *Rmst*) of isocortex^23,65^. Whether transcriptionally related cells in isocortex and subiculum also show similar connectivity and functional properties remains unclear.

Functionally cells in layers 5 and 6 of isocortex that project to the thalamus are classified as drivers or modulators, respectively, based on their synaptic properties^66–69^. Layer 5 cortico-thalamic neurons provide input to higher order thalamic nuclei, via collaterals of subcortical projections^70^. These are powerful, large and depressing, “driver” inputs, often sufficient to induce action potentials (AP) for relay to other cortical areas^71–73^. These projections are thought to not form synapses in RT^71,74^. In contrast layer 6 cortico-thalamic neurons form smaller and more distal inputs onto thalamic neurons to evoke small and facilitating EPSCs and metabotropic responses with stronger stimulation. They are called “modulators” because they influence how postsynaptic neurons process other inputs. These neurons form collaterals to the inhibitory RT^68,75,76^.

The functional properties of subicular inputs to the thalamus are not known. We investigated characteristics of PV+ subiculo-thalamic projections by assessing distinguishing features of the driver-modulator classification for corticothalamic neurons. We report that PV+ projections may form axon collaterals within the subiculum but not in the RT. In the AV they evoke small, graded EPSCs, which were able to elicit firing when postsynaptic cells were close to threshold, reminiscent of modulators feature. However, they act only via ionotropic receptors and show response depression similar to drivers. Morphologically we found that PV+ subiculo-thalamic neurons make synapses on proximal dendrites and somata similar to drivers but with relatively small terminal varicosities (0.8 to 2 µm^2^ cross section area), which is more common for modulators^48,77^ (but see^66^). Accordingly, we suggest that PV+ subiculo-thalamic neurons cannot unequivocally be classified according to functional criteria applied to cortico-thalamic neurons of layers 5 and 6. Our investigation of the synaptic properties of PV+ subiculo-mammillary projections have only been exploratory but suggest that similar to subiculo-thalamic PV+ neurons, these neurons show small ionotropic receptor-dependent EPSCs and synaptic depression (see Supplementary Fig. 6).

An unresolved finding of the current study is that in situ hybridization indicated that the vast majority of subicular PV+ neurons expressed VGAT and that consistently VGAT was co-expressed with VGLUT1 or VGLUT2 in a fraction of subicular cells, presumably including those which expressed PV. In agreement with the detection of VGAT transcripts we also found VGAT immunoreactivity in terminals of PV+ projections in AV and mMB. However, we found no indication of GABAergic transmission after stimulation of PV+ terminals in AV and mMB (Fig. 3, Supplementary Fig. 6,7). This incongruity has also been reported for the mossy fibers in the dentate gyrus^78^, which mediate excitatory responses in CA3 but contain low levels of VGAT. In case of mossy fibers it has been suggested that GABA release may be transient during development^79,80^ or after periods of enhanced excitability^78^. In mossy fibers transient GABA co-transmission coincides with up-regulation of the machinery for GABA synthesis. In GABAergic interneurons, PV+ expression levels change with neuronal activity^81,82^, which in turn can regulate the GABA production machinery^83^. Subsequent studies will have to explore if and under which circumstances an activity dependent switch in transmitter release can occur in these cells.

Regarding electrophysiological phenotype and neuronal morphology, we were able to show that deviating from most neighbouring pyramidal cells, PV+ principal cells in the dorsal subiculum are regular spiking and possess straight and unbranching apical dendrites. Although we recorded from PV+ pyramidal cells in deep and more superficial layers, this is certain only for AV targeting neurons in deep layers as MB projecting PV+ cells were not specifically traced. Whereas the functional significance of the firing phenotype of subicular neurons -bursting or regular firing -is not known, it is thought that these biophysical properties are related to local and distant connectivity as well as specific patterns of network activity^18,24,84^. Similarly, the relevance of the specific dendritic morphology is not known but likely relates to the composition of synaptic inputs to these neurons^85^.

Location, dendritic morphology, out-put specificity and functional properties thus suggest that the two populations of PV+ pyramidal neurons in the dorsal subiculum are tied into specific streams of information. The nature of this information remains to be studied but based on their connectivity with AV and MB may be related to theta oscillatory networks for memory formation and to the processing and transfer of spatial information^39,40,55,62,86,87^. Although AV and MB, likely play specific roles in memory formation and spatial processing, these two structures themselves are densely connected and work together during spatial and mnemonic computations^87–89^. A similar situation has been described for two population of L5 PV+ neurons in the motor cortex^22^, which display different levels of PV expression, different laminar distribution and different downstream connectivity but show complementary cooperation in motor control. It is thus possible that also the two populations of PV+ projection neurons in the subiculum provide functionally related information to separate components of a cooperative AV-MB network.

Taken together and independent of the precise content of transmitted information, we show that PV is expressed in two separate populations of excitatory neurons of the distal subiculum. These neurons show a high degree of target selectivity and present with specific functional and morphological features suggesting that PV-expression in archicortex like in isocortex identifies subgroups of excitatory neurons with specific connectivity and functional properties.

## Acknowledgments

We thank Kira Balueva for guidance with viral vectors and in situ hybridizations. Our work was supported by the Deutsche Forschungsgemeinschaft (DFG) FOR 2143, RTG 2154 and SFB 1461 (Project-ID 434434223) to P.W and DFG FOR 2143 and CRC-TRR384 to A.K.

## Author contributions

G.B. and P.W. devised the study. G.B., A.F. and P.W. designed experiments. A.F. performed virus-and dye-based anatomical tracing, histological analysis of synapses, of in situ hybridizations and PV expression and prepared samples for electron microscopy. G.B. performed and analyzed: anatomical tracings, electro-physiological experiments, neuronal reconstructions, and associated immunohistochemistry. N.W. and A.K. performed electron microscopy. K.A.E. and C.W. analyzed transcriptomic data. K.K. performed histological experiments, J.M. helped with confocal microscopy of synapses. G.B. directed the study and wrote the first draft of the manuscript. G.B. and P.W. wrote the final version of the manuscript with the help of all authors.

## Methods

### Animals

All procedures involving experimental animals were in accordance with the German Animal Welfare Act and approved by the local authorities. PV-Cre mice^8^ were purchased from Jackson laboratories (Repository number 008069) and maintained as heterozygous colonies or crossed with Ai9 Cre reporter mice^90^ (Jackson laboratories, Repository number 007909). C57BL6 mice were purchased from Janvier Labs, RRID:MGI:2670020. All experimental procedures were carried out on heterozygous PV-Cre or PV-Cre::Ai9 mice or on C57bl/6 mice between 4 to 8 weeks. Mice were maintained in a 12 h light-dark cycle under standard group housing conditions and were provided with food and water ad libitum.

### Recombinant AAV vectors

AAV-FLEX-mCherry (containing mCherry fused to DREADD inverted in a flip-excision (FLEX) cassette), AAV-FLEX-GFP, AAV-CaMKII-GFP (containing EGFP-T2A-FlpO under the CaMKII promoter), AAV-DIO-ChR2-YFP (AAV1/2-EF1a-DIO-hChR2(E123T/T159C)-EYFP-WPRE-pA, addgene #35509) were produced as described previously^10,91^. Briefly, virions containing a 1:1 ratio type 1 and type 2 capsid proteins were produced by transfecting human embryonic kidney (HEK) 293 cells with the respective rAAV backbone plasmids along with AAV1 (pH21), AAV2 (pRV1) and adenovirus helper plasmid pFdelta6 using the calcium phosphate method. 48 hours post transfection, cells were harvested and rAAVs were purified using 1 mL HiTrap heparin columns (Sigma) and concentrated using Amicon Ultra centrifugal filter devices (Millipore). Infectious rAAV particles (viral titer) were calculated by serially infecting HEK293 cells stably expressing Cre-recombinase and counting GFP-positive cells. AAVs were used at a titer of 1x10^7^ to 1x10^8^ infectious particles per ml. AAV-Dlx-ChR2-YFP (serotype 5) was a gift from Ilka Diester and used at a titer of 1,6*10^13^ vg/mL.

### Stereotaxic surgeries

Stereotaxic surgeries were performed as described^91^. For Analgesia mice were pre-treated with Carprofen (Rimadyl 5 mg/kg) and then anesthetized with 3% isoflurane in O2 by inhalation. Anaesthesia was maintained on 1.5–2% isoflurane throughout surgery. Mice were fixed with their heads in a stereotaxic frame (Kopf Instruments, USA) and body temperature was maintained by a heating mat placed underneath the animal. For local anaesthesia we subcutaneously injected the skin above the skull with prilocaine (Xylonest 2%). After exposure of the skull, small holes were drilled relative to Bregma. Stereotaxic coordinates for the subiculum were: AP -2.8, ML -0.97, DV -1.75 for anatomy and AP -2.78, ML ±1.1, DV -1.75 for optogenetic experiments. Stereotaxic coordinates for the AV and MB were: AP -0.57/-0.85 mm, ML -2.5/-2.65 mm, DV -3.55, angle 30° for AV and AP -2.7 mm, ML -0.1 mm, DV -5.25 for MB. rAAV was injected at a volume of 0.3-0.5 µl using a Hamilton microliter syringe (Hamilton Company, USA) over a period of 5 min. Fast Blue (Polysciences, cat number 17740-1), 0.5% Red IX Retrobeads or Green Retrobeads (both Lumafluor Inc.) were injected at a volume of 0.3 µL. The syringe was retracted after a 10 min interval. Burr holes were filled with bone wax, the skin was replaced and fixed with Vetbond tissue adhesive (3M, USA). Analgesic treatment was given as needed. During the recovery period mice were housed individually. Mice were used for immunohistochemistry or electrophysiology 21-28 days after rAAV injection and 4-7 days after injection of retrograde tracers.

### Immunohistochemistry

Mice were deeply anaesthetized with pentobarbital (Narcorene 50 mg per 30 g body weight i.p.) and transcardially perfused with phosphate buffered saline (PBS, pH 7.4) for 4 minutes followed by 4% paraformaldehyde (PFA) in PBS for 10 min. Brains were post-fixed in PFA 4% for at least 4 h at 4°C, embedded in 4% agar in PBS and cut with a Leica VT1200S vibratome (section thickness: 50µm).

Free-floating sections were permeabilized for 30 min in 0.4% Triton X-100 in PBS and blocked for 30 min in PBS containing 0.02% NaN_3_, 4% normal goat serum (NGS), 0.2% Triton X-100 at room temperature. Primary antibodies were diluted to working concentrations in PBS containing, 0.1% Triton X-100, 2% NGS and incubated with sections for 24 to 48 h at 4°C. Primary antibodies used were: rabbit polyclonal anti-GFP (1:1000, A6455, Invitrogen, California, USA), mouse monoclonal anti GFP (1:1000, A11120, Invitrogen, California, USA) rabbit polyclonal anti-PV (1:1000 to 1:2000, PV25, Swant, Switzerland), mouse monoclonal anti-PV (1:5000, PV235, Swant, Switzerland), rabbit polyclonal anti-RFP (1:2000, 600-401-379, Rockland, Pennsylvania, USA), mouse monoclonal anti-RFP (1:2000, 200-301-379, Rockland, Pennsylvania, USA), rabbit polyclonal anti-VGAT (1:1000, 131003, Synaptic Systems, Germany), guinea pig polyclonal anti-VGLUT2 (1:5000, AB2251, Millipore, Germany), guinea pig polyclonal anti-VGLUT2 (1:500, 135-404, Synaptic Systems, Germany). Following primary antibody incubation, sections were washed three times for 10 minutes in PBS with 1% NGS at room temperature and incubated with secondary antibodies for 2-3 hours at room temperature. Secondary antibodies used were: Goat anti-mouse Alexa Fluor 488 (1:1000, A11001, Invitrogen, California, USA), goat anti-rabbit Alexa Fluor 488 (1:1000, A11008, Invitrogen, California, USA), goat anti-rabbit Cy3 (1:1000, 111-165-144, Jackson Immunoresearch, Pennsylvania, USA), goat anti-mouse Cy3 (1:1000, 115-166-003, Jackson Immunoresearch, Pennsylvania, USA), goat anti-mouse Cy5 (1:1000, 115-175-146, Jackson Immunoresearch, Pennsylvania, USA), goat anti-guinea pig Cy5 (1:500, 106-175-003, Jackson Immunoresearch, Pennsylvania, USA). Sections were then washed in PBS containing 1% NGS and twice in PBS alone for 10 minutes. After a quick rinse in distilled water, sections were mounted onto glass slides (Roth, Germany) and cover-slipped using Mowiol (Sigma, Massachusetts, United States).

Brain slices that were processed for immunohistochemistry after patch clamp-recordings were post-fixed in 4% PFA at 4°C for 24 h. Sections were then washed 3x in PBS, treated with a permeabilizing and blocking solution (0.5% Triton, 10% NGS, 0.05% NaN_3_ in PBS) for 5 h at RT. The primary antibody solution contained 0.05% NaN_3_ in PBS. Primary anti-bodies used were mouse monoclonal anti-GFP (1:0000, A11120, Invitrogen, California, USA) and guinea-pig polyclonal anti-VGLUT2 (1:500, 135-404, Synaptic systems, Germany). Primary antibodies were applied for 2 days at 4°C. The secondary antibody solution contained 0.05% NaN_3_ in PBS. Secondary antibodies were goat anti-mouse Alexa Fluor 488 (1:1000, A11001, Invitrogen, California, USA) and goat anti-guinea pig CY5, (1:500, 106-175-003, Jackson Immunoresearch, Pennsylvania, USA). These were applied for 24 h at 4°C. For visualization of biotin-filled cells we used streptavidin Alexa 488, 555 or 594 (1:500, S11223, S21381 or S11227, Invitrogen, California, USA) for 48 h at 4°C. Sections were mounted on slides (Roth, Germany) and cover-slipped using mounting medium (Vectashield, Vectorlab, Germany).

### In situ hybridization

Fresh unfixed mouse brains were frozen on dry ice and stored at -80°C until further processing. Coronal sections with a thickness of 20 µm were prepared using a cryostat (Leica; CM3050) and collected on Polysine Adhesion Slides (Thermo Fisher Scientific, Waltham, USA). RNA probe hybridization and subsequent washes were performed as described previously^92^. Probes were synthesised using fluorescein RNA labelling mix (Roche Diagnostics, Mannheim; Germany) or digoxygenin (DIG) labelling mix (Roche Diagnostics, Mannheim, Germany) and detected with peroxidase-conjugated anti-fluorescein or anti-digoxygenin antibodies. Peroxidase activity was detected with Cy3-or FITC-tyramide conjugates. Fluorescein-and Cy3-tyramide conjugates were synthesised as described previously^93^. After labelling, peroxidase activity was blocked by quenching with 100 mM glycine-HCl (pH 2.0) solution containing 0.1% Tween. Subsequent steps were performed as in Webster et al. (2020)^94^. Sections were counterstained with 4, 6-diamidino-2-phenylindole (DAPI) (Sigma-Aldrich, Munich, Germany).

PV mRNA was detected using a mixture of 3 probes: 1) 217 bp of Exon 4, primers AGGTTCTGCCTGTGACCTTG and AAGCTTTGACAGCCGCATAC; 2) 248 bp of Exon 3, primers CCCTCTCCCCTGTCCTTCTTT and ATGGGAACTTTGGGTGCTATC; 3) 573 bp of Exon 5 including 3’ UTR, primers CCTCCACTCTGGTGGCTGAA and TTTCTCTTTTCAGGTATTTTATCACA. For VGAT a 927 bp probe of exon 2 was generated. Primers were GCTACCTGGGGTTGTTCCTCA and CGAAGTGTGGCACGTAGATG. For VGLUT2 a 962 bp probe was generated of exon 12, primers CTATTCGTTGGACCCATCACC and CGGCCGCAGAAATTGCAATCCCCAAAC. VGLUT1 mRNA was detected with a 719 bp probe (sequence was identical to the probe RP_050310_01_B09 from the Allen Mouse Brain Atlas dataset, mouse.brain-map.org/experiment/show/69014470), primers CAGAGCCGGAGGAGATGA and TTCCCTCAGAAACGCTGG.

DNA templates for probe synthesis were PCR-amplified from C57Bl6 mouse genomic DNA. Amplified DNA fragments were cloned into the pBSK backbone, which was linearised and transcribed with T3 RNA polymerase using DIG RNA Labeling Mix orT7 RNA polymerase using Fluorescein RNA Labeling Mix according to the manufacturer protocol (all from Roche Diagnostics, Mannheim, Germany).

### Microscopy and image analysis

Images were acquired with either a fluorescent microscope (Axio-Imager M2 with Apotome, Carl Zeiss, Germany) or a confocal microscope (LSM 880 confocal laser scanning microscope with Airyscan, Carl Zeiss) using a Plan-Apochromat 10x objective (NA 0.45, Zeiss), a Plan-Apochromat 20x objective (NA 0.8, Zeiss), an EC Plan-Neofluar 20x/0.50 M27 or a Plan-Neofluar 40x oil-immersion objective (NA 1.4, Zeiss). Zen 2.3 was used to merge images from different channels.

#### In situ imaging and graphical representation

Images were acquired with a Zeiss Axio-Imager M2 with Apotome, Zeiss, 20x objective. Outputs were tiled stack images of 6 optical slices with 5 µm in total. Colocalization analysis was done by manual counting. Results were represented in heat maps where the dorsal subiculum was divided into 12 subfields along the proximal/distal axis and superficial/deep layers and the density of double positive over total single positive cells within each subfield was plotted.

#### Analysis of synapses and labelling intensities

For synapse analysis Airyscan confocal images, 40x objective, with a field of view of about 300x300µm and stacks of 100 to 150 optical slices (interval 0.1-0.2 μm) were captured. Scaling per voxel was 0.1 x 0.1 x 0.1-0.2 μm. Synaptic puncta on fluorescently labelled axons were analysed using Imaris software (BitPlane). A solid surface best matching the filament anatomy and demarcating the postsynaptic domain was generated using the “surface” tool. Background was subtracted. The “smoothing” tool was disabled, to avoid artificial uniformity. GFP voxel histograms were used to determine the threshold for background exclusion. Channels for presynaptic markers were filtered based on the spatial relationship with the neuronal surface (removal of presynaptic fluorescent signals outside the surface, and postsynaptic fluorescent signals inside the surface). For quantification of presynaptic puncta, the minimum diameter was set to 0.6 µm as measured in “slice view” mode^82^. Synapses were quantified as positive puncta over the total fiber volume reconstructed in the field of view.

For analysis of PV labelling intensities, images were acquired at a Zeiss LSM800 confocal microscope with a Plan-Neofluar 40x oil-immersion objective (NA 1.4, Zeiss). Stacks of 100 to 130 optical slices with an interval of 0.3 μm were captured. Analysis of PV expression levels was carried out by generating three-dimensional isosurfaces for each labeled soma (number of voxels >1) and PV immunofluorescence intensities were quantified automatically as a mean of all voxels in arbitrary units (Imaris 8.0.0, Bitplane AG)^81,82^. Settings remained the same during image acquisition for all samples.

#### Anatomical mapping and morphological analysis of biocytin filled cells

To map the location of recorded neurons, images of the entire slice containing the biocytin filled cells were taken at an Axio-Imager M2 with Apotome a using a Plan-Apochromat 10x objective. The AV or MB were identified and the filled cells were registered in the respective atlas plate (Paxinos and Franklin, 2012), according to the spatial relationship with regional borders. For recordings in the subiculum, filled cells were reported on a single representative plate using the end of CA1 (proximal-distal axis) and the pia border (the deep-superficial axis) as reference.

For reconstruction of biocytin filled cells, Airyscan confocal images were acquired using a Plan-Apochromat 20x objective (NA 0.8, Zeiss) or a Plan-Neofluar 40x oil-immersion objective (NA 1.4, Zeiss) and a pixel size of 0.1 x 0.1 x 0.180, or 0.04 x 0.04 x 0.180 µm (x,y,z). Cells were reconstructed using Imaris 9 (Bitplane, SouthWindsor, CT, USA) using the z-stack fluorescent images generated from Airyscan-confocal microscopy. A detailed surface rendering of Alexa Fluor 555-Biocytin filled neurons, VGLUT2-immunoreactive presynaptic terminals (CY5-secondary antibody) and ChR2-YFP labelled projections (Alexa Fluor 488 secondary antibody) was created. The neuronal surface was used to demarcate postsynaptic and presynaptic domains. Neuronal surfaces were created using the “create surface” tool with the “smoothing” tool disabled to avoid artificial uniformity of the cell surface. “Background subtraction” was selected to detect fluorophore in a volume containing the soma, the proximal dendrites and distal dendrites. The average diameter of the smallest dendrites, measured with the line tool in “slice view”, was taken to determine the minimum diameter setting. The threshold was selected on the histogram of AF555-Voxel to include as much of the neuron as possible while excluding any background. Proximal dendrites were defined as the primary and secondary branches of the dendrites. To reconstruct presynaptic signal and YFP+ fibers the staining within the filled neuron was filtered out using the masking function. The minimum diameter was set to 0.6 µm^95^. The “region growing” function was selected to identify surfaces of different size and the “morphological split” function to detect the borders. The sensitivity for detecting puncta was adjusted using the histogram based on quality of the signal to detect puncta without creating artefacts. Using the objects statistic algorithm, we first located the total number of surfaces adjacent to the filled neuron within a distance of 0.5 µm, thus representing the putative synaptic contacts. Then we co-localized VGLUT2+ surface with YFP+ surfaces using the algorithm “overlapping volume ratio” set above 5%” which takes into account a minimum of two optical sections. The number of putative YFP-and VGLUT2-positive contacts per surface area of the filled cell was calculated to compare densites of putative synapses on proximal dendrites and soma to distal dendrites. The diameter size in Supplementary Figure 12 was measured on random distal and proximal dendrites of each reconstructed cell.

The size of presynaptic terminals was quantified (Cross section area µm^2^) using Fiji software^96^. A maximum intensity projection of 12 optical slices for each filled thalamic cell (n=3) was chosen randomly. Only the green channel (YFP signal) was selected, smoothed and thresholded. Terminals were detected as particles with size ≧ 0.35 µm and circularity ≧ 0.3^66^.

### Electron microscopy

A total of 3 adult mice was used in the present study. Animals were deeply anesthetized by pentobarbital (Narcoren 50 mg per 30 g body weight i.p.), and the hearts were surgically exposed for transcardial perfusion fixation using a fixative solution containing 4% PFA, 0.05% glutaraldehyde and15% (v/v) saturated picric acid as described previously^97^.

For double immunoelectron microscopy, sections were prepared as described previously^97^. Briefly, sections (60 µm), containing the antero-ventral nucleus of the thalamus, were blocked, and then incubated in a mixture of primary antibodies (GFP: rabbit, 1:3000, Abcam; VGLUT2: Guinea pig, 1:300, Frontiers Institute) diluted in 50 mM Tris-buffered saline (TBS) containing 3% Normal goat serum (NGS). Subsequently, the sections were incubated in a mixture of secondary antibodies: biotinylated goat anti-rabbit antibody (1:100, Vector Laboratories) and goat anti-guinea pig (Fab fragment 1:100) coupled to 1.4 nm gold particle (Nanoprobes, Stony Brook, NY) made up in TBS containing 1% NGS. After washes in TBS, sections were washed in double-distilled water, followed by silver enhancement of the gold particles with an HQ silver kit (Nanoprobes). Subsequently, the sections were incubated in ABC (Vector Laboratories). Peroxidase was visualized with DAB (0.05% in Tris buffer (TB) using 0.01% H_2_O_2_ as substrate. The sections were treated with 1% OsO4 and then contrasted in 1% uranyl acetate. They were dehydrated in a series of ethanol and propylene oxide and flat embedded in epoxy resin (Durcupan ACM, Sigma, UK). After polymerization, short series of ultrathin sections were cut at 70 nm thickness using an ultramicrotome (Reichert Ultracut E; Leica, Vienna, Austria) and analysed in a JEOL JEM-2100 Plus electron microscope.

Cross section area of GFP+ boutons containing VGLUT2 and the diameter size of contacted dendrites was measured and calculated on images of random planes of boutons and dendrites.

### Electrophysiology

#### Slice preparation and solutions

Brain slice preparation, storage and recordings were performed as described before^98^. Three to four weeks after stereotaxic injections PV-Cre or PV-Cre::Ai9 mice were briefly anesthetized with isoflurane and the brain was extracted. Acute 300 µm-thick sagittal or coronal brain slices were prepared on a sliding vibratome (HM 650 V, Thermofisher, USA) in ice-cold oxygenated NMDG-based solution (containing in mM: 92 NMDG, 2.5 KCl, 1.25 NaH_2_PO_4_, 30 NaHCO_3_, 20 HEPES, 25 glucose, 5 Na-ascorbate, 4 Na-pyruvate, 0.5 CaCl_2_·2H_2_O, and 10 MgSO_4_·7H_2_O; pH 7.4). Slices were kept for 10 min in the same solution at 35°C before being transferred to a recovery solution (containing in mM: 92 NaCl, 2.5 KCl, 1.25 NaH_2_PO_4_, 30 NaHCO_3_, 20 HEPES, 25 glucose, 5 Na-ascorbate, 4 Na-pyruvate, 2 CaCl_2_·2 H2O, and 2 MgSO_4_·7 H_2_O; pH 7.4) at room temperature for at least 30 min. Slices were placed in the recording chamber under an upright microscope (Olympus BX50WI, 20x water-immersion objective, Volketswil, Switzerland) and continuously perfused at room temperature with oxygenated ACSF (containing in mM: 122 NaCl, 2.5 KCl, 1.25 NaH_2_PO_4_, 24 NaHCO_3_, 12.5 D(+)-glucose, 2 CaCl_2_ and 2 MgSO_4_, pH 7.4).

Cells were visualized through differential interference contrast optics. Infrared images were acquired with a CCD Camera. For targeted recordings cells were identified by the respective fluorescent signal before and after patching. Cells were patched using borosilicate glass pipettes (TW150F-4) (World Precision Instruments, Sarasota, USA) pulled with a P1000 horizontal puller (World Precision Instruments, Sarasota, USA) to a final resistance of 3.5 – 5 MΩ. A K^+^-based intracellular solution (containing in mM: 135 KGluconate, 10 HEPES, 3 KCl, 1 MgCl_2_, 0.2 EGTA, 6 Na_2_-phosphocreatine, 3 Mg-ATP, 0.3 Na-GTP; pH 7.3, 290 – 305 mOsm) was supplemented with 1 mg/ml neurobiotin (Vector Labs, Servion, Switzerland) and used for measurements of active and passive properties and all current clamp recordings. A Cs^+^-based intracellular solution (containing in mM: 125 CsMetSO_3_, 2 MgCl_2_, 10 HEPES, 1 EGTA, 2 CsCl, 10 Na_2_phosphocreatine, 2 MgCl_2_, 4 Mg-ATP, 0.3 Na_2_-GTP, 5 QX314-Cl, supplemented with 1 mg/ml neurobiotin; pH 7.3, 290–305mOsm) was used in all voltage-clamp protocols except for the tonic current measurement which was measured with a CsCl intracellular solution (containing in mM: 130CsCl, 10 HEPES, 1 EGTA, 10 CsOH, 8 NaCl,2 Mg-ATP, 0.3 Na-GTP, 5 QX314-Cl, supplemented with 1 mg/ml of neurobiotin; pH 7.3, 290–305mOsm). For these solutions, liquid junction potential was measured and subtracted from the recordings. Signals were amplified using a HEKA USB10 amplifier and filtered at 3 kHz and digitized at 10 kHz.

#### Recording protocols, optogenetic stimulation and analysis

For all recordings, immediately after gaining whole-cell access, cell resistance (Rm), holding current (pA) and cell capacitance (Cm) were measured in voltage-clamp at -65 mV while applying 100 ms-long, 5-10 mV hyperpolarizing steps (5 steps average). Then the recording was switched to current-clamp to measure the resting membrane potential (RMP).

Additional measurements of active and passive properties of subicular and thalamic neurons were performed in current-clamp. To measure sag ratio, we applied squared somatic current injections (-50 to -300 pA for 1 s, 3 injections/cell) to hyperpolarize neurons below -100 mV from membrane potentials between -60 to -65 mV. The sag ratio was defined as the ratio of the steady state voltage (average voltage from 400-500 ms) relative to baseline, divided by the minimum voltage (usually occurring within 100 ms of the onset of the hyperpolarizing step) relative to baseline. The same protocol used for sag ratio induced repetitive single burst discharge in thalamic neurons and was used to measure the number of action potential per single rebound burst. Squared current injections of increasing amplitude (step size, 10 to 20 pA, 800-1000 ms) were used to depolarize subicular or thalamic neurons. Action potential properties were measured at the rheobase or 50 pA above rheobase^57^ as indicated in Supplementary Fig. 10. Rheobase was defined as a current injection level equal to the first intensity able to trigger firing. We measured: spike threshold (defined as membrane potential where dV/dt exceeded 10 V/s^99^); spike amplitude (difference between threshold and action potential peak); full width at half maximum voltage (FWHM, elapsed time between the voltage crossings at half maximal amplitude during the rising and falling phase); maximal spike rising and maximal spike falling (measured on the first derivative of the action potential (dV/dt)); afterdepolarization (ADP, calculated for each spike by finding peak voltage following the downstroke of the action potential relative to baseline^100^); frequency (total number of spikes generated by 1 s depolarization). We measured the instantaneous firing for the first two spikes of the response^57^.

Whole-field blue LED (Cairn Res, Faversham, UK) (470 nm, duration: 1 ms, power: 20-30 %) was used to stimulate ChR2-YFP-expressing fibers during voltage-clamp recordings (-70 mV and 10 mV) of AV and MB neurons. The LED light optical power at different percentages of maximum was measured with an optical spectrometer placed under the objective (20x, NA 1, water immersion, Olympus). To calculate irradiance (mW/mm^2^) we calculated the surface area (mm^2^) as the circle area using the diameter in mm obtained by dividing the objective Field number (22) by its magnification (20X). 8% power thus corresponded to 1.7 mW/mm^2^ and 30% corresponded to 5.6 mW/mm^2^.

To test for excitatory and inhibitory currents, using Cs^+^-based intracellular solution, we recorded EPSCs at -70 mV and subsequently depolarised the membrane potential of the cell to +10 mV to record IPSCs. Post synaptic currents were elicited through single light pulses every 10-15 s, with a -10 mV hyperpolarizing step to control for the access resistance. The latency from LED onset, EPSC half-width and EPSC decay time constant were measured using Clampfit 10.2 (Molecular Devices, USA). After a stable baseline of > 5 min, drugs were applied into the bath: 10 µM NBQX, 100 µM D, L-APV, 10 µM SR95531 or Gabazine (GBZ), 1 µM TTX (all HelloBio, Ireland), 100 µM 4-AP (Sigma, Massachusetts, USA), 100 µM LY367385 (Tocris, UK).

Paired light stimulations at 1, 3, 5, 6.6, 10 and 20 Hz were used to assess short-term plasticity. Three to six responses were elicited for each frequency, with an interval of 15 s between each protocol. EPSC amplitudes were measured on the averaged trace in Clampfit10.2 (Molecular Devices, USA). Response ratios were calculated by dividing the amplitude of the n-th EPSC by the amplitude of the first EPSC.

For extended testing for GABA_A_ receptor-mediated responses, cells were held at + 10 mV with a Cs-based internal solution and ChR2-YFP-expressing afferents were light stimulated with 10 stimuli at 20, 40 or 100 Hz. To test for tonic inhibition, fibres were stimulated at 5 Hz for 4 min while cells were held at -65 mV in a CsCl-based internal solution. Baseline values of holding current were measured as mean during the last 3 min before stimulation. The steady state holding current was measured 10 min after the end of stimulation as average of the last 3 min. The effect on the holding current was measured as the difference between steady state value from baseline. To test for postsynaptic GABA_B_-receptor-mediated responses, subicular ChR2-YFP-expressing afferents were stimulated with 20 light pulses delivered every 30 s at 20, 40, 100 or 200 Hz while cells were held at -50 mV in voltage-clamp using a K-gluconate based intracellular solution. Per stimulation frequency 3-5 traces were recorded and averaged.

### Transcriptomic data analysis

Transcriptome analysis was performed on SMARTseq counts data by Ding et al. (2020)^23^ [https://data.nemoarchive.org/biccn/grant/u19_zeng/zeng/transcriptome/scell/10x_v2/mouse/ processed/analysis/RHP/] using RStudio (RStudio version 1.4.1106, R version 4.1.0) and the Seurat package (version 4.3.0).

First, raw counts data were normalized to CPM values (counts per million) and cells were categorized into PV-positive and PV-negative subgroups using a CPM threshold = 5, corresponding to ∼20% and 80% of the total cell population. PV-positive cells were next filtered and re-clustered. To this end, PV-positive cells were first log-normalized, screened for highly variable genes (settings: selection.method = ’vst’, clip.max = log2(#cells), nfeatures = 2000), and rescaled, followed by principal component and a cluster analysis. For cluster visualization, the first two t-SNE dimensions were extracted. Additionally, heatmaps were computed to compare clusters in the PV-positive subgroup on differentially expressed genes as well as pre-selected genes of interest.

### Statistical analysis

Sample sizes are given in the figure legends or text. “n” represents either mice or neurons as indicated. Statistical analysis was performed using MATLAB and GraphPad Prism version 6. We used two-tailed unpaired Student’s t test, multiple unpaired t-test and Holm Sidák multiple comparison test as appropriate. When n represented multiple neurons from a relatively low number of subjects, i.e. Figure 5 C,D, we accounted for intra-class correlation due to clustered data in form of random effects. Accordingly, unless stated otherwise in the text, we applied a linear mixed-effects model using MATLAB (y = Xb+Zu+ε, where y is the response variable, X is the predictor variable, b is a vector of the fixed-effects regression coefficients, Z is the random effect variable, u is a vector of the random effects and ε is a vector of the residuals). This approach maintains information about variability and avoids under-estimations of the p value. For Zscore normalisation we used the formula Z = (x−µ)/σ, where x is the observed value, µ is the mean of the sample and theta is the standard deviation of the sample. A p value of less than 0.05 was considered significant. Unless stated otherwise, in figures bars with error bars refer to means ± SEM.

## Data and code availability

This paper does not report original code.

Any additional information required to reanalyze the data reported in this paper is available from the lead contact upon request.

