## Supplementary Figure 1-12 and Table 1 for "Parvalbumin expression identifies subicular principal cells with high projection specificity"

This file contains

Supplementary Figures 1 to 12

Supplementary Tables 1

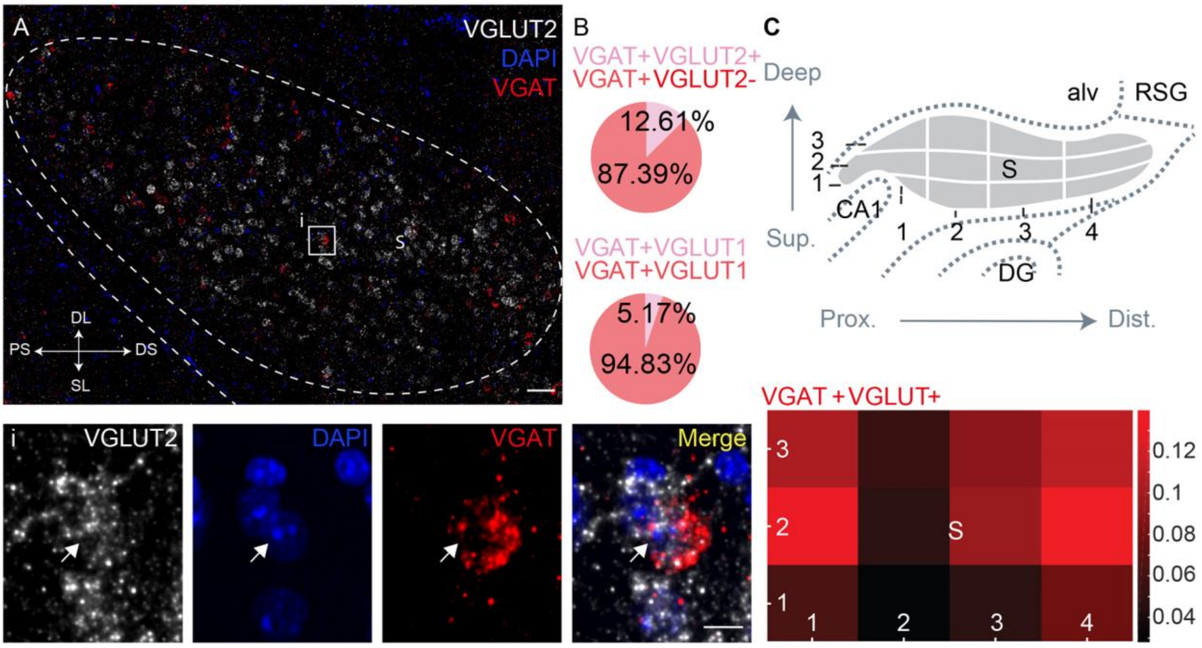


**Figure S1. A subgroup of VGAT expressing neurons in the subiculum also expresses VGLUT1 and VGLUT2.** A) Double fluorescent in situ hybridization on a coronal brain section showing co-localization of VGLUT2 (white) and VGAT (red) mRNA in the subiculum. Nuclei are stained with DAPI (blue). Scale bar 50 µm. i Magnification of the boxed area in A. The arrow indicates a cell positive for both VGAT and VGLUT2 mRNA. Scale bar 10 µm. B) A small percentage of subicular VGAT+ neurons co-expressed VGLUT2 (n=817 cells in 2 mice) or VGLUT1 (n=774 cells in 1 mouse) mRNA. C) To describe the location of VGLUT1/2 expressing VGAT+ neurons, we divided the subiculum into 12 sub-fields along the proximal/distal axis and superficial/deep layers (upper panel). The relative occurrence of double labelled neurons is depicted in a corresponding heat map (lower panel). CA1, cornus ammonis 1, DG, dentate gyrus, S, subiculum; alv, alveus; RSG, retrosplenial cortex granular.

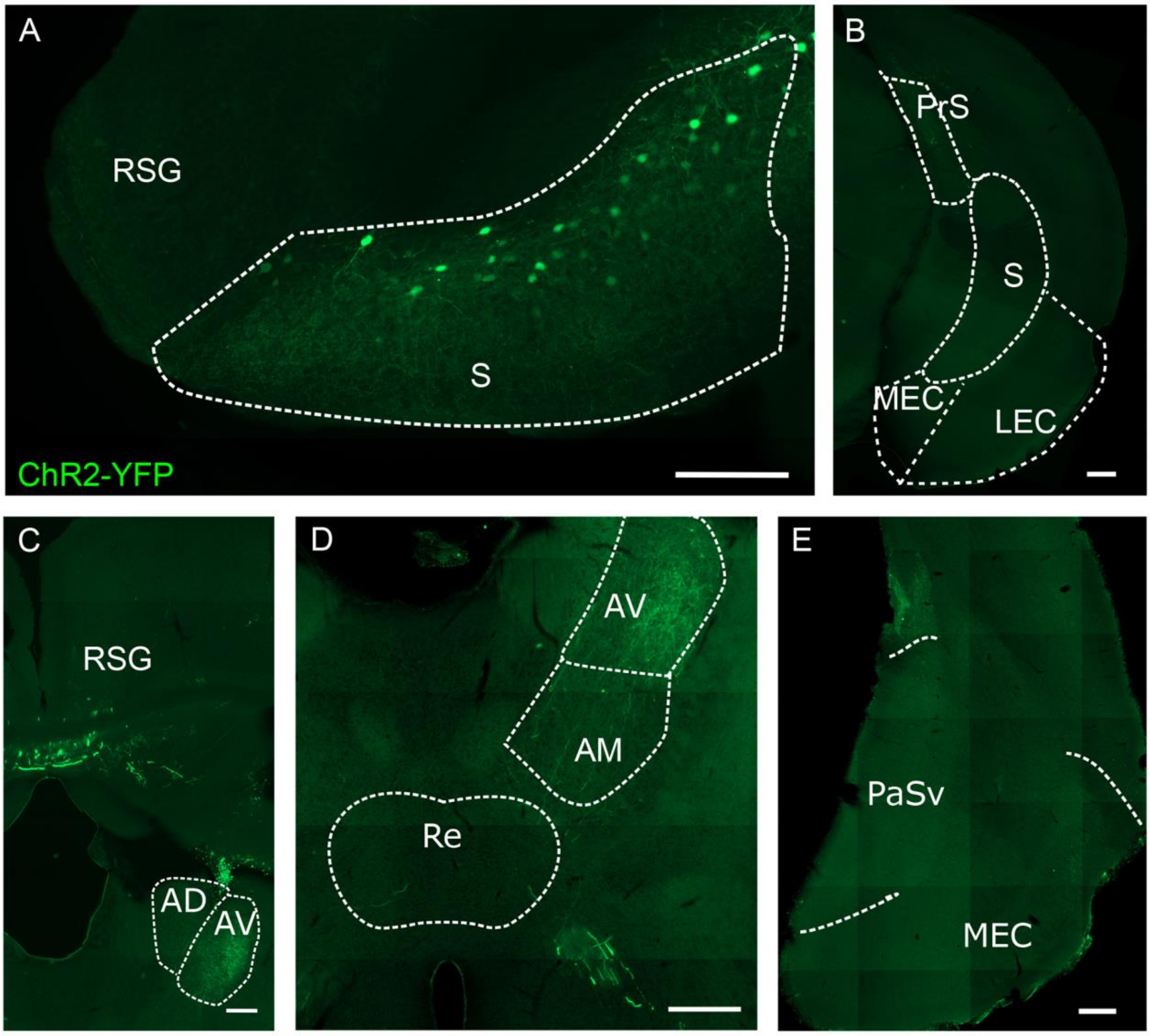


**Figure S2. Parvalbumin positive projections from the subiculum spare many of the known efferent areas of the subiculum**. A) Site of injection of a Cre-dependent AAV expressing ChR2-eYFP in the dorsal subiculum (S) of a PV-Cre mouse at Bregma level -2.80 mm. B) Fibres were absent from the retrosplenial cortex (RSG), nucleus reuniens (Re), presubiculum (PrS), antero-dorsal and antero-medial nucleus (AD, AM), medial as well as lateral enthorinal cortex (MEC, LEC) and ventral parasubiculum (PaSv).

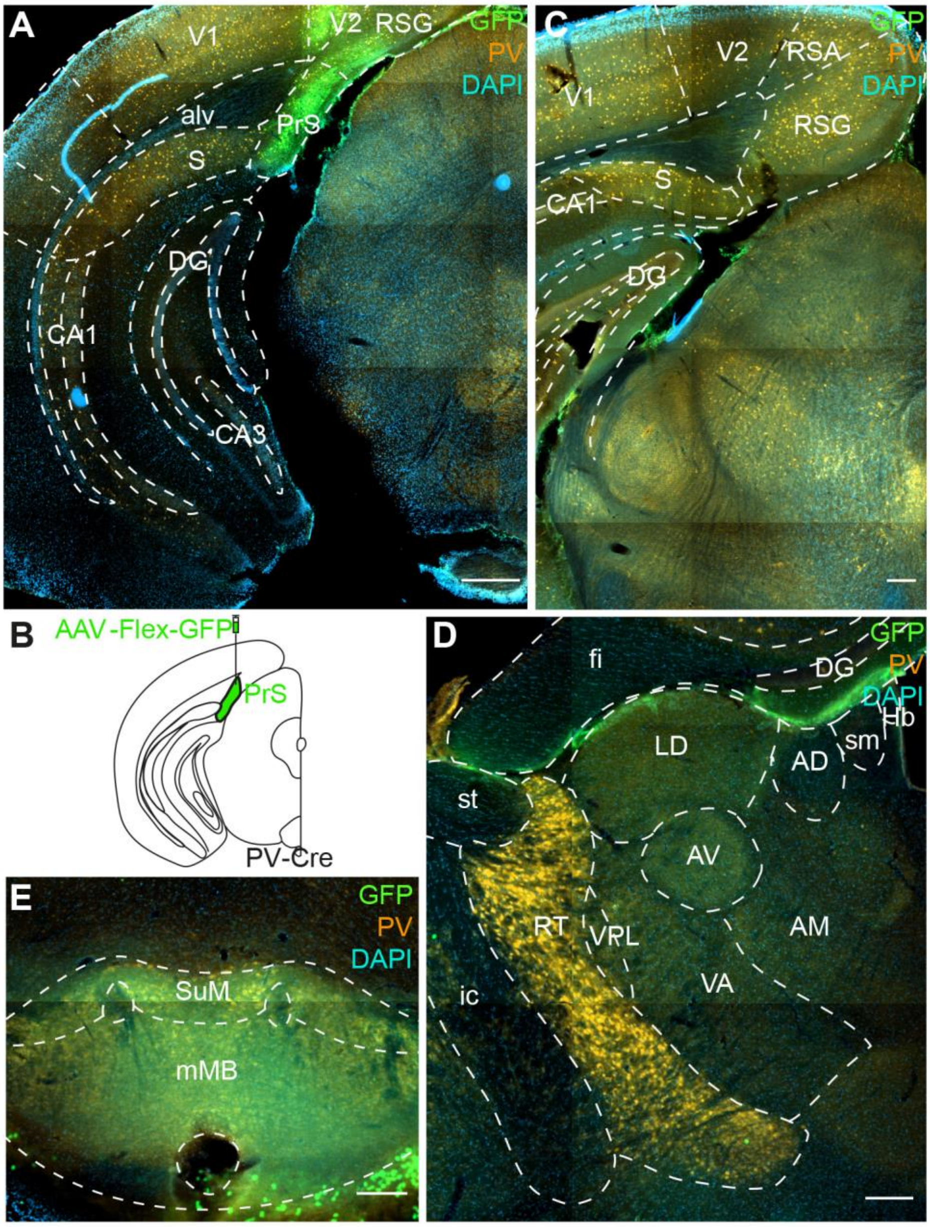


**Figure S3. PV-positive neurons of the pre-subiculum do not project to the anteroventral thalamus or mammillary bodies.** A) Coronal brain section of a PV-Cre mouse, injected with a Cre-dependent AAV expressing GFP in the pre-subiculum (PrS) as illustrated in B. The section was immuno-stained for GFP (green), PV (orange) and DAPI (blue). The transduced area was mainly centered in the presubiculum but included also parts of the secondary visual (V2) and retrosplenial (RSA, RSG) cortices. Scale bar 500 µm. C) The subiculum (S) is completely free of viral infection. Scale bar 200 µm. D) Coronal section including the thalamic area of the mouse in A. No projecting fibers were detectable in any of the thalamic nuclei. Scale bar 200 µm. E) Coronal section including the hypothalamic area of the mouse in A. No GFP-positive fibers were visible in the supramammillary bodies (SuM) or in the medial mammillary bodies (mMB). Scale bar 200 µm. AD, antero-dorsal nucleus of the thalamus, alv, alveus, AM, antero-medial nucleus of the thalamus, AV, antero-ventral nucleus of the thalamus, CA, cornu ammonis, DG, dentate gyrus, fi, fimbria, ic, internal capsule, LD lateral dorsal nucleus of the thalamus, Hb, medial habenular nucleus, RT, reticular thalamus, sm, stria medullaris of the thalamus, st, stria terminalis, V1, primary visual cortex, VA, ventral anterior thalamic nucleus, VPL, ventral posterolateral thalamic nucleus.

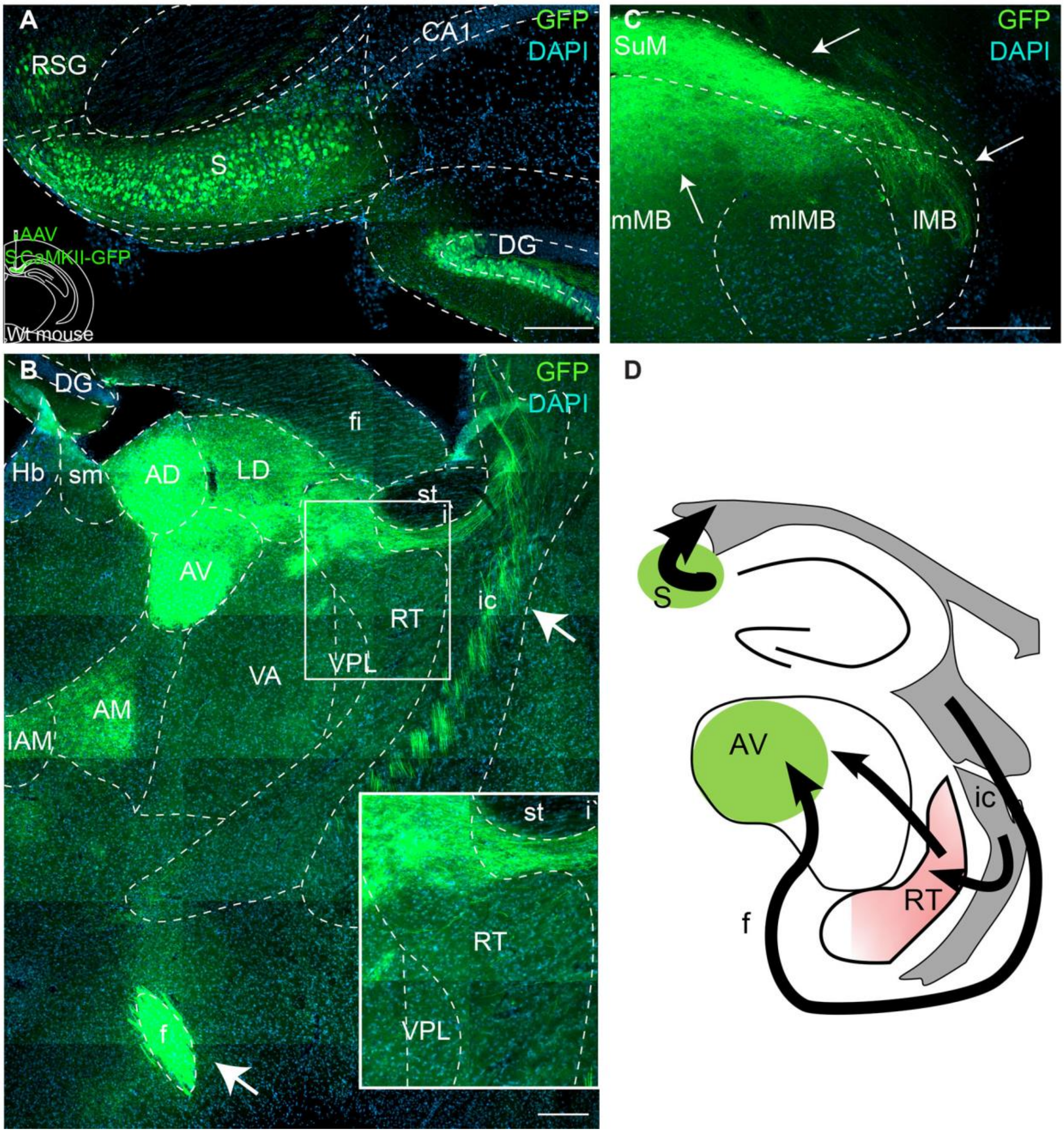


**Figure S4. Transduction of subicular principal cells shows projections through fornix and internal capsule broadly targeting nuclei of the anterior thalamus and mammillary bodies.** A) Coronal brain section of a wild type (Wt) mouse injected with an AAV expressing GFP under the control of the CaMKII promoter, showing broad transduction of subicular (S) principal cells with some viral spread to retrosplenial cortex (RSG) and dentate gyrus (DG). Scale bar 200 µm. B) Dense GFP-positive fiber projections are visible within both internal capsule (ic) and fornix (f) (white arrows), targeting anterior and latero-dorsal nuclei of the thalamus (LD). Note the occurrence of labeled fibers also in the reticular thalamus (RT, inset shows close up of boxed area). Scale bar 200 µm. C) GFP-positive fiber projections can be found in different subregions of the mammillary bodies (mMB, medial, mlMB, mediolateral, lMB, lateral mamillary body, white arrows). D) Simplified scheme illustrating possible subicular projection pathways to the thalamus. The projection passing through the internal capsule may also synapse with cells of the reticular nucleus, which in turn projects to the thalamus. AD, antero-dorsal nucleus of the thalamus, AM, antero-medial nucleus of the thalamus, AV, antero-ventral nucleus of the thalamus, CA, cornu ammonis, fi, fimbria, IAM, inter-antero-medial thalamic nucleus, LD lateral dorsal nucleus of the thalamus, Hb, medial habenular nucleus, RT, reticular thalamus, sm, stria medullaris of the thalamus, st, stria terminalis, SuM, Supramamillary body, VA, ventral anterior thalamic nucleus, VPL, ventral postero-lateral thalamic nucleus.

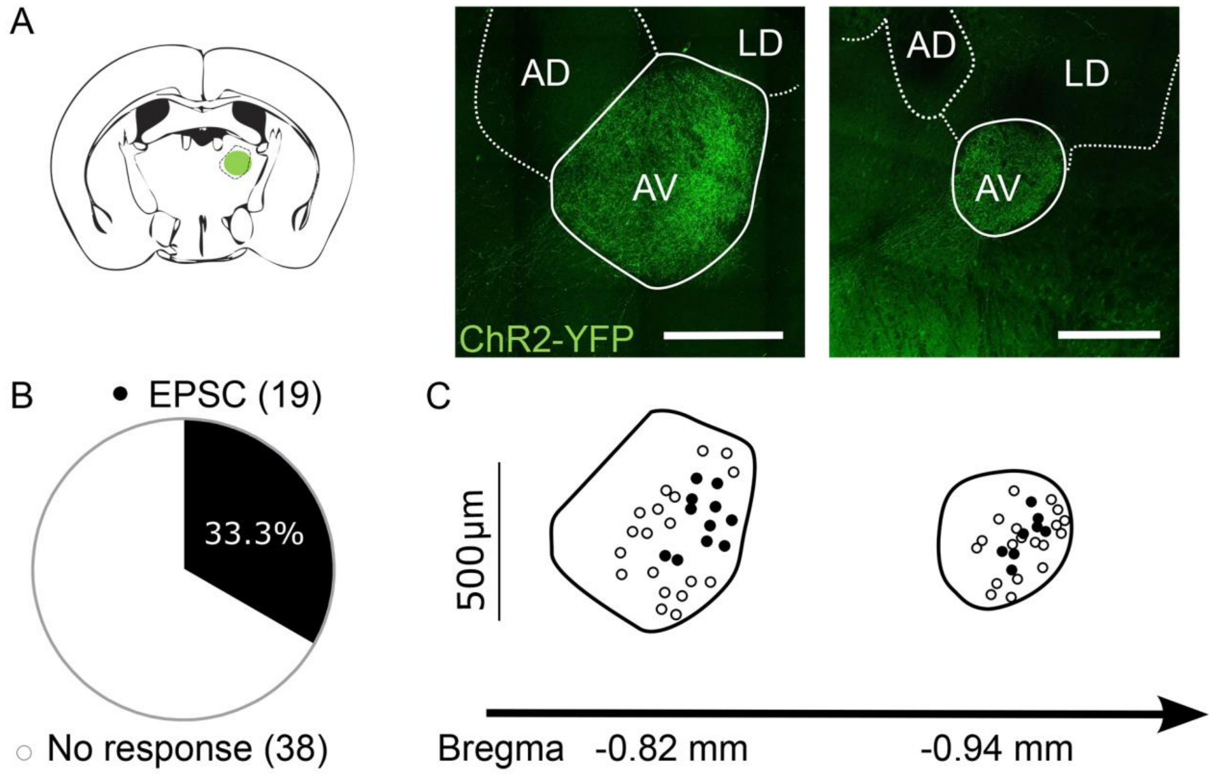


**Figure S5. Topographical distribution of responsive neurons in coronal slices of the anteroventral thalamus.** A) Left, cartoon of a coronal plate illustrating the location of PV+ terminals from the subiculum in the AV (green circle). Right, confocal image of a coronal slice showing the location of the ChR2-YFP positive fibres (green) in the lateral part of the dorso-caudal AV. B) 33% of recorded AV neurons showed excitatory responses after the light stimulus, C) Whole-cell recorded neurons were filled with neurobiotin and registered onto coronal planes of the AV (black dots, EPSC; white dots, no response). Scale bar, 500 µm. AV, antero-ventral nucleus, AD, antero-dorsal nucleus, LD, latero-dorsal nucleus of the thalamus.

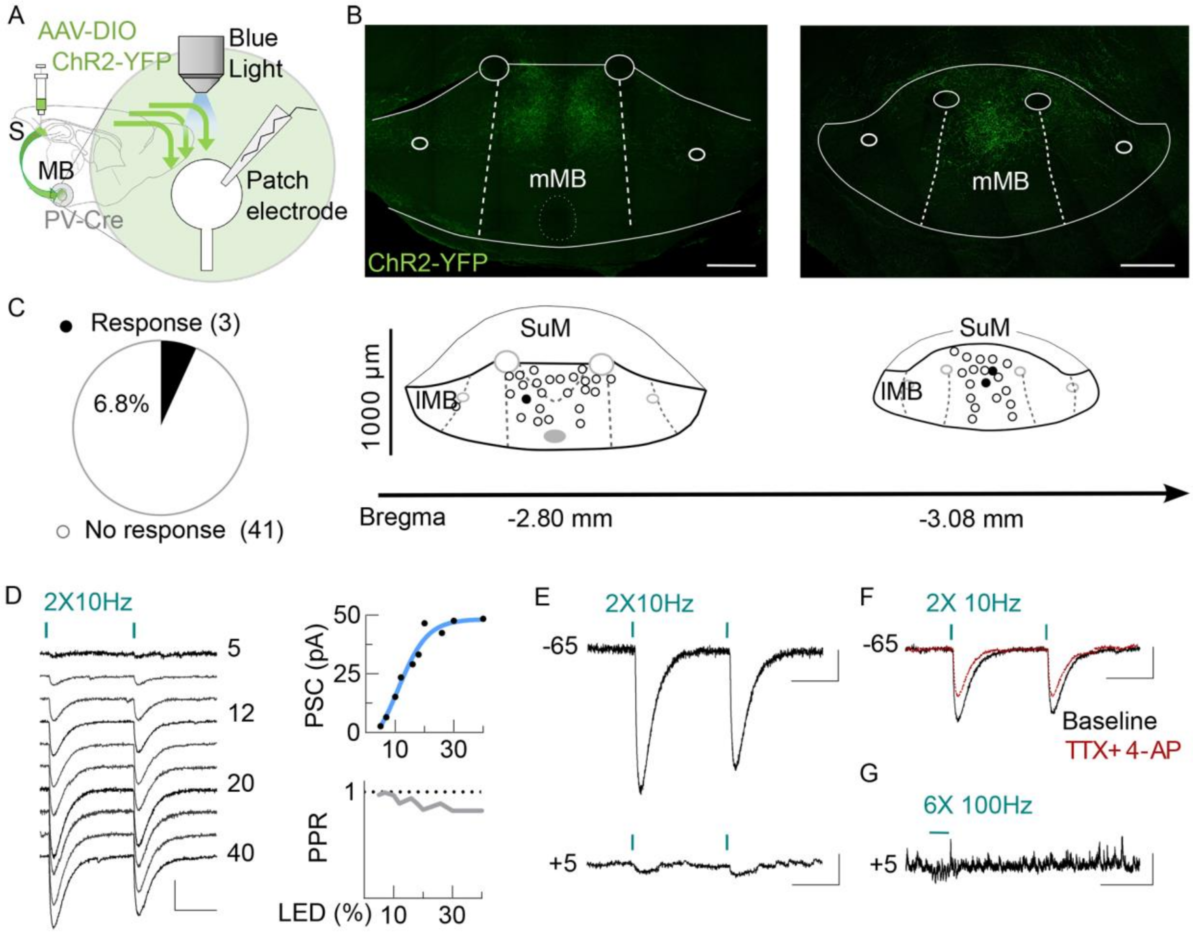


**Figure S6. PV+ projections from the subiculum form functional excitatory synapses in the medial mammillary body.** A) Cartoon illustrating the experimental setup to record post-synaptic responses in neurons of the mammillary body (MB) during photo stimulation of PV+ ChR2-YFP expressing fibres (green) from the subiculum (S). B) Top, confocal images of coronal section showing the location of ChR2-YFP-expressing fibers (green) in the medial MB (mMB). Bottom, recorded neurons were filled with neurobiotin and registered in coronal planes of the MB (black dots, response (EPSC) and white dots, no response). Scale bar, 250 µm. lMB, lateral mammillary body, SuM, supramammillary body. C) Pie chart showing the percentage of responsive neurons. D) Left, examples of current responses of a mammillary body neuron to photo stimulation of PV+ subicular afferents at different intensities. Right, input–output curve (top) and paired pulse ratio (bottom) as a function of light intensity. Scale bar 20 pA, 50 ms. E) Example of a cell showing EPSCs (top) but no IPSCs (bottom). Scale bars 20 pA, 100 ms. F) . Light stimulated responses are recovered by 4-AP in the presence of TTX. Scale bar 20pA, 100ms (LED 15% or 3 mW/mm$^2$). G) High frequency stimulation did not evoke IPSCs. Scale bar 20 pA, 1 s (LED 30%).

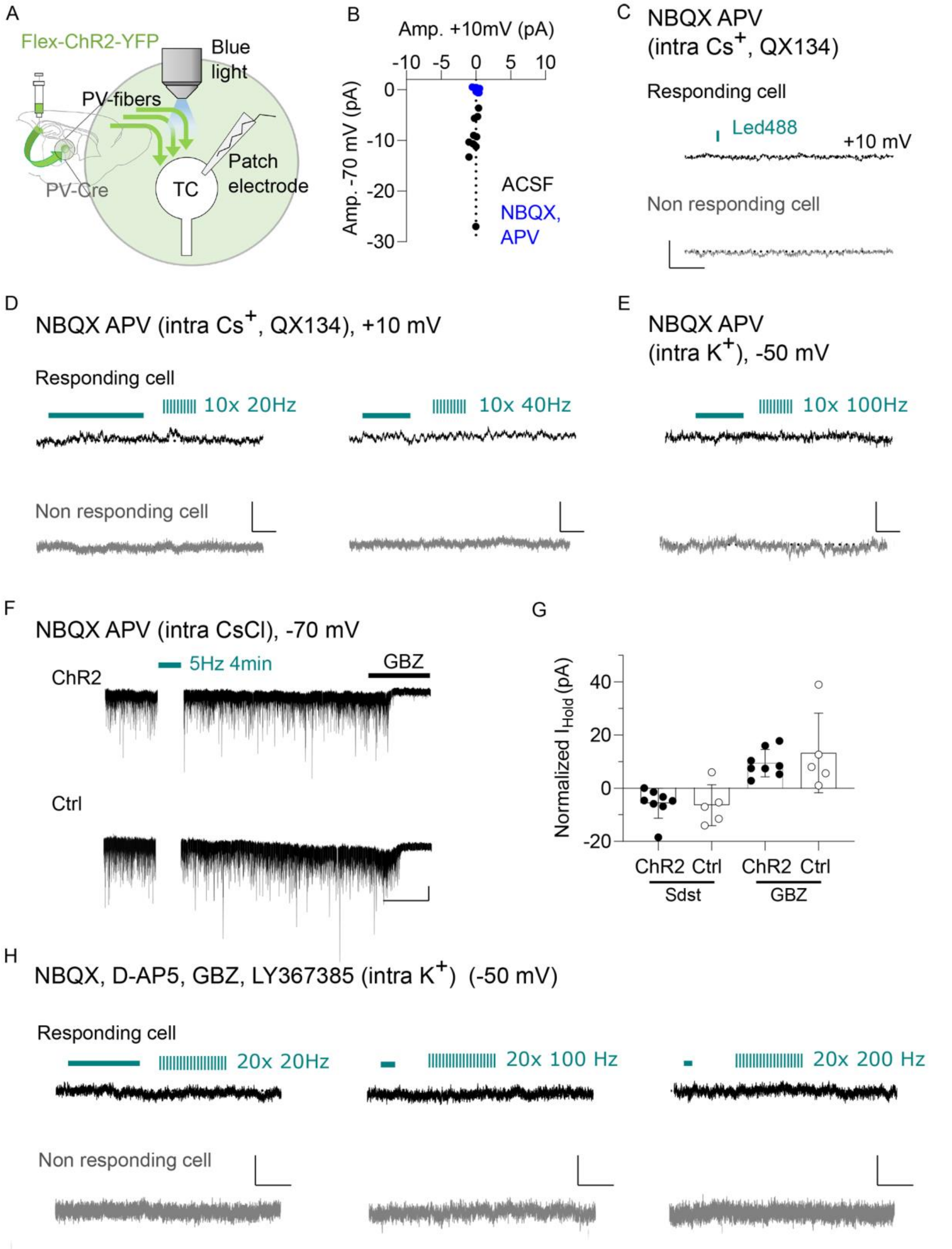


**Figure S7. Photo stimulation of PV+ subicular projections with different protocols does not elicit GABAergic responses in the AV.** A) Cartoon illustrating the experimental configuration. Whole-cell patch-clamp recording of a thalamic cell (TC) during optogenetic stimulation of PV+ fibers from the subiculum. B) Measurement of current amplitude at 10 mV and -70 mV of n=10 cells before and after pharmacological antagonism of glutamatergic ionotropic receptors with NBQX and APV. C) Example traces of a cell that responded with EPSCs (responding cell) (n=10) and a cell that showed no EPSCs (non-responding cell) (n=14)

tested for IPSCs in the presence of NBQX and APV (Cs+MetSO4 based pipette solution) (scale 5 pA, 50 ms). D,E) Trains of light stimuli at different frequencies did not activate a detectable GABA-A receptor-mediated current, neither in cells that showed light evoked EPSCs nor in cells that showed no light evoked EPSCs. D) N=4 cells per type at 20 Hz; n=5 cells per type at 40 Hz. Cs+MetSO4 based pipette solution; scale 5pA,100ms. E) N=5 cells per type at 100 Hz. $K^+$ gluconate based pipette solution; scale 5 pA, 50 ms. F) Voltage clamp recording of the holding current at -70 mV before, during and after a long optogenetic stimulation at 5 Hz of PV+ terminals expressing ChR2 or GFP (Ctrl). GABAergic currents are directed inward (CsCl based pipette solution). The steady state (Sdst) is the difference between the value of the holding current at baseline and the plateau value at 10 min post-stimulation (measured over 3 min). GBZ is the difference between the plateau value and the value after Gabazine application. G) The light-stimulation of ChR2+ terminals did not increase the change in holding current beyond the average value obtained with the stimulation of control terminals. ChR2 (n=8), Ctrl (n=5). Gabazine induced a comparable change in holding current independent of whether terminals expressed GFP or ChR2. H) Trains of 20 stimuli at different frequencies in the presence of a cocktail of antagonists did not elicit a response mediated by GABA-B receptors in cells with EPSC responses (responding cell) or cells without EPSC responses (non-responding cell) (3 mice, n=5 responding and n=5 non-responding cells). K+ gluconate-based pipette solution; scale 5 pA, 500 ms.

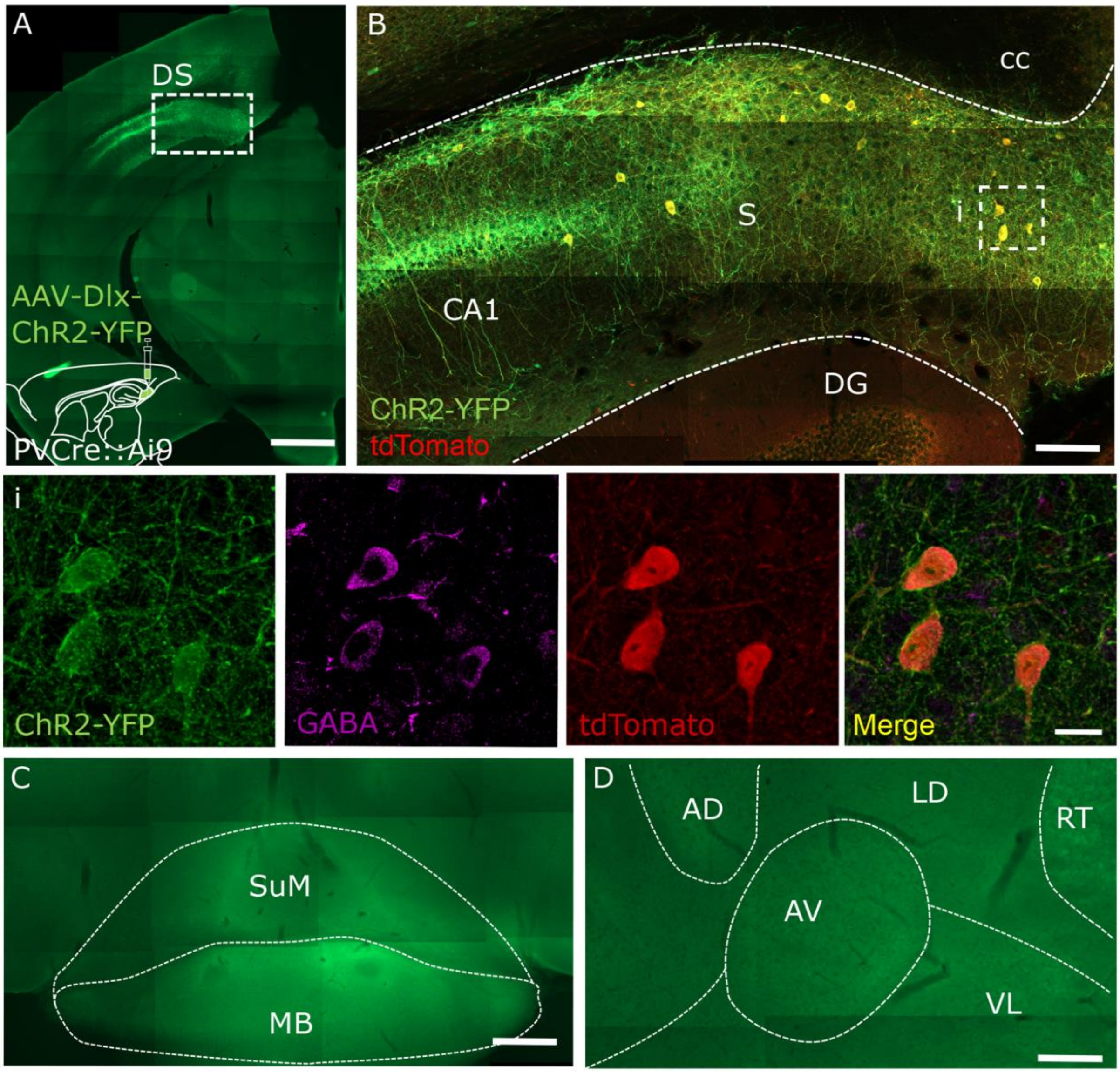


**Figure S8. Dlx-expressing PV+ neurons in the subiculum are not projecting to AV and MB.** A) Injection of AAV-Dlx-ChR2-YFP into the dorsal subiculum (DS) of PV-Cre::Ai9 mice (n=3).Scale bar 500 µm. B) ChR2-YFP is expressed in tdTomato-positive cells of the dorsal subiculum. Scale bar 200 µm. b1) Magnification of the boxed area in B showing three neurons immunoreactive for ChR2-YFP, GABA and tdTomato. Scale bar 20 µm. The mammillary body (MB) (C) and the antero-ventral nucleus (AV) (D) are devoid of YFP positive fibers. Scale bar 200µm. AD, antero-dorsal nucleus, CA1, cornu ammonis region 1, cc, corpus callosum, DG, dentate gyrus, LD, latero-dorsal nucleus, RT, reticular nucleus, S, subiculum, SuM, supramammillary body, VL, ventro-lateral nucleus.

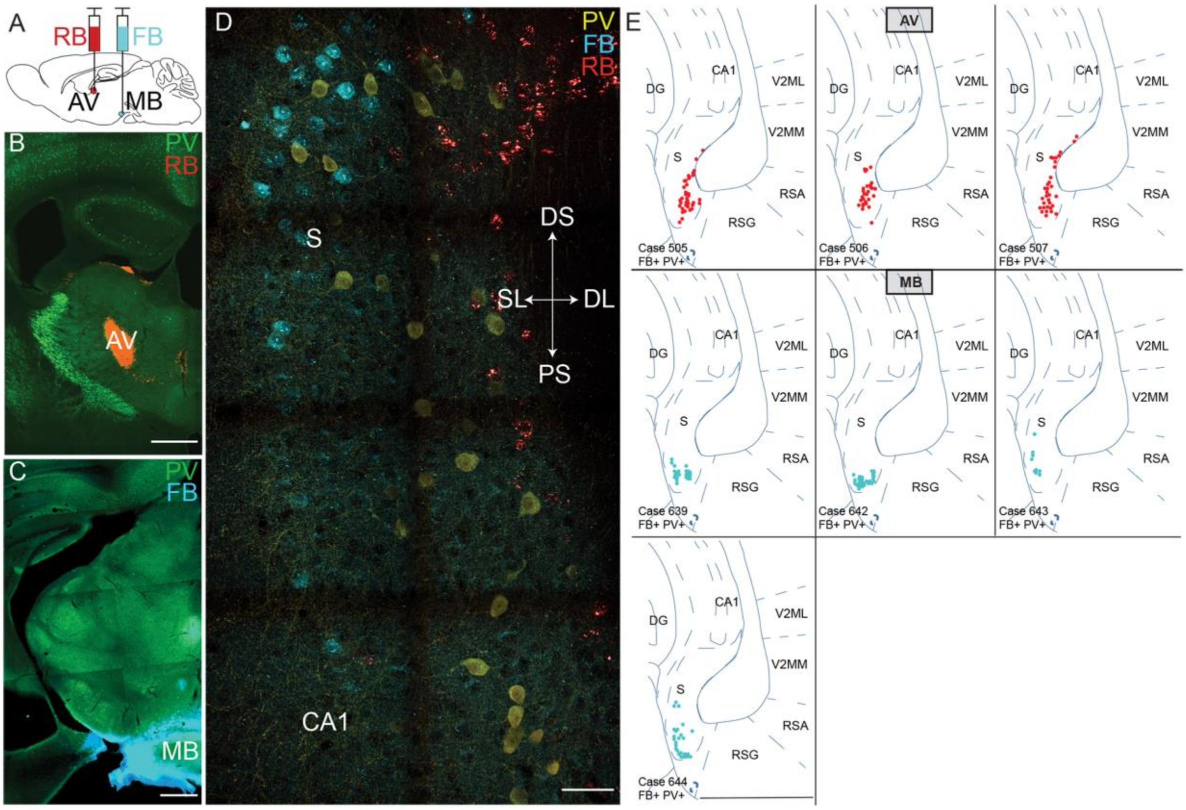


**Figure S9. Retrograde tracing of projections to the anteroventral thalamus and the mammillary body labels separate populations of PV+ neurons in the subiculum.** A) Illustration of the injection sites of the retrograde tracers red retrobeads (RB) and fast blue (FB) in the antero-ventral thalamus (AV) and mammillary body (MB), respectively. B) Brain section showing the spread of red retrobeads in the AV (red fluorescence). PV immunoreactivity is shown in green. Scale bar 500 µm. C) Brain section showing the spread of fast blue in the MB (blue fluorescence) and immunostaining for PV in green. Scale bar 500 µm. D) Same animal as in A and B. Retrogradely labelled cells in the distal subiculum are spatially separated with cells traced from the AV and the MB in deep and superficial layers, respectively (PV immunostaining in yellow). Scale bar 50 µm. E) Additional retrograde tracing experiments were done with single injections of fast blue into either the AV or the MB. The three panels on the top show the location of retrogradely labelled PV+ cells after injection of FB into the AV. The four panels on the bottom show the location of retrogradely labelled PV-positive cells after injection of FB into the MB. Each panel is one animal, summarizing 3 consecutive brain sections. Each dot is one cell. AV, antero-ventral nucleus of the thalamus; CA, cornu ammonis; DG, dentate gyrus; DL, deep layer; DS, distal subiculum; FB, fast blue; MB, mammillary nucleus; PS, proximal subiculum; PV, parvalbumin; RB, red retrobeads; RSA, retrosplenial cortex agranular; RSG, retrosplenial cortex granular; S, subiculum; SL, superficial layer; V2ML secondary visual cortex, mediolateral area, V2MM, secondary visual cortex, medio-medial.

A

| | tdTom+ Bead+ PC | | | | tdTom+ PC | | | | tdTom- PC | | | | |
|---|---|---|---|---|---|---|---|---|---|---|---|---|---|
| **Subthreshold** | **Mean** | | **s.e.m.** | **n** | **Mean** | | **s.e.m.** | **n** | **Mean** | | **s.e.m.** | **n** | |
| Resting membrane potential (mV) | -61.16 | ± | 1.30 | 8 | -61.25 | ± | 0.65 | 21 | -65.84 | ± | 0.83 | 5 | |
| $R_N$ (MΩ) | 445.51 | ± | 34.61 | 8 | 448.03 | ± | 34.22 | 12 | 158.60 | ± | 13.25 | 5 | *** |
| Sag ratio | 0.82 | ± | 0.02 | 8 | 0.73 | ± | 0.02 | 14 | 0.67 | ± | 0.02 | 5 | |
| **Suprathresold, at rheobase** | | | | | | | | | | | | 5 | |
| Spike threshold (mV) | -42.20 | ± | 1.72 | 6 | -45.85 | ± | 0.99 | 18 | -46.72 | ± | 22.05 | 5 | |
| Spike Amplitude (mV) | 81.24 | ± | 4.22 | 7 | 83.96 | ± | 2.17 | 18 | 95.93 | ± | 0.49 | 5 | * |
| Max spike rising dV/dt (mV/ms) | 217.66 | ± | 34.88 | 6 | 225.13 | ± | 17.54 | 18 | 425.93 | ± | 1.17 | 4 | *** |
| Max spike falling dV/dt (mV/ms) | -64.34 | ± | 8.00 | 6 | -63.39 | ± | 4.41 | 18 | -95.54 | ± | 25.49 | 4 | |
| Full-width at half-maximum (FWHM) (ms) | 1.37 | ± | 0.19 | 7 | 1.32 | ± | 0.06 | 18 | 1.01 | ± | 12.60 | 5 | * |
| IF of 1st and 2nd spike (Hz) | 12.01 | ± | 6.64 | 7 | 10.91 | ± | 1.98 | 18 | 154.60 | ± | 0.03 | 5 | *** |
| Afterdepolarization (mV) | 0.64 | ± | 0.24 | 7 | 3.99 | ± | 1.65 | 10 | 6.50 | ± | 0.72 | 5 | |
| **Suprathreshold, at rheobase+50pA** | | | | | | | | | | | | | |
| Frequency (Hz) | 23.67 | ± | 2.76 | 6 | 26.58 | ± | 0.91 | 19 | 23.85 | ± | | 2 | |
| IF of 1st and 2nd spike (Hz) | 59.23 | ± | 9.66 | 7 | 69.35 | ± | 4.38 | 19 | 160.68 | ± | 0.15 | 5 | *** |
| 2nd Spike threshold (mV) | -38.71 | ± | 1.04 | 7 | -38.97 | ± | 0.78 | 19 | -39.45 | ± | 8.97 | 5 | |
| Amplitude 1st spike (mV) | 83.02 | ± | 4.66 | 7 | 88.73 | ± | 2.20 | 19 | 95.58 | ± | 0.81 | 5 | |
| Amplitude 2nd spike (mV) | 70.35 | ± | 2.35 | 7 | 69.46 | ± | 2.34 | 19 | 73.19 | ± | 1.27 | 5 | |
| Max 2nd spike falling dV/dt (mV/ms) | -53.71 | ± | 5.56 | 7 | -51.57 | ± | 3.63 | 19 | -50.24 | ± | 2.77 | 5 | |
| Max 2nd spike rising dV/dt (mV/ms) | 161.67 | ± | 17.82 | 7 | 150.97 | ± | 11.19 | 19 | 225.34 | ± | 3.45 | 5 | *** |
| 1st spike FWHM (ms) | 1.24 | ± | 0.12 | 7 | 1.17 | ± | 0.06 | 19 | 1.01 | ± | 30.52 | 5 | |
| 2nd spike FWHM (ms) | 1.39 | ± | 0.12 | 7 | 1.35 | ± | 0.08 | 19 | 1.14 | ± | 0.05 | 5 | |

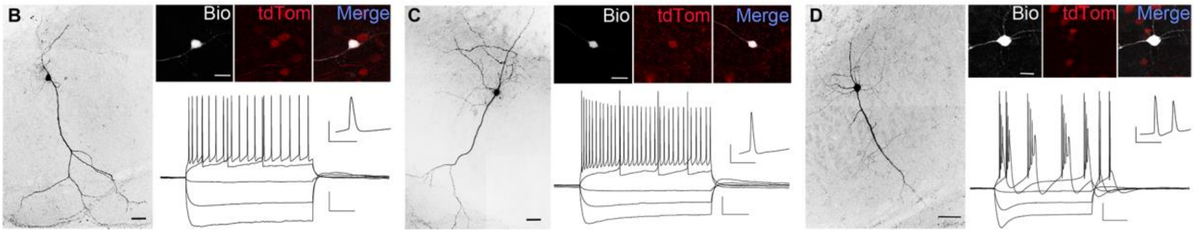


**Figure S10. The electro-physiological properties of PV+ excitatory neurons differ from neighboring principal cells in the distal subiculum**. A) Table summarizing passive and active properties of tdTomato+ pyramidal cell like neurons, containing beads, tdTomato+ pyramidal cell like neurons in mice without retrobead injections and neighboring tdTomato- pyramidal cells. Holm-Sidak´s multiple comparison test *** p<0.001, Unpaired t-test * p<0.05 between tdTomato+ and tdTomato-PCs. B) Example of a biocytin filled tdTomato+ pyramidal cell like neuron, containing beads (scale bar 50 µm) and its voltage response. Scale 20 mV, 200 ms. Inset shows the first action potential of the trace at rheo-base +50 pA at higher temporal resolution. Scale 40mV, 10ms. Top Right shows an image of the same cell with biocytin and tdTomato colocalization. Scale bar 20 µm. C) Same as B for a tdTomato+ pyramidal cell like neuron in a mouse that was not injected with retrobeads. D) Same as B and C for a tdTomato-negative pyramidal cell in the distal subiculum, where biocytin and tdTomato do not colocalize. Note the similarity in electro-physiological properties and morphology between tdTomato+ pyramidal cell like neurons with and without beads, which are regular firing and have apical dendrites almost devoid of oblique dendrites. In contrast tdTomato-negative pyramidal cells are burst firing and have oblique dendrites.

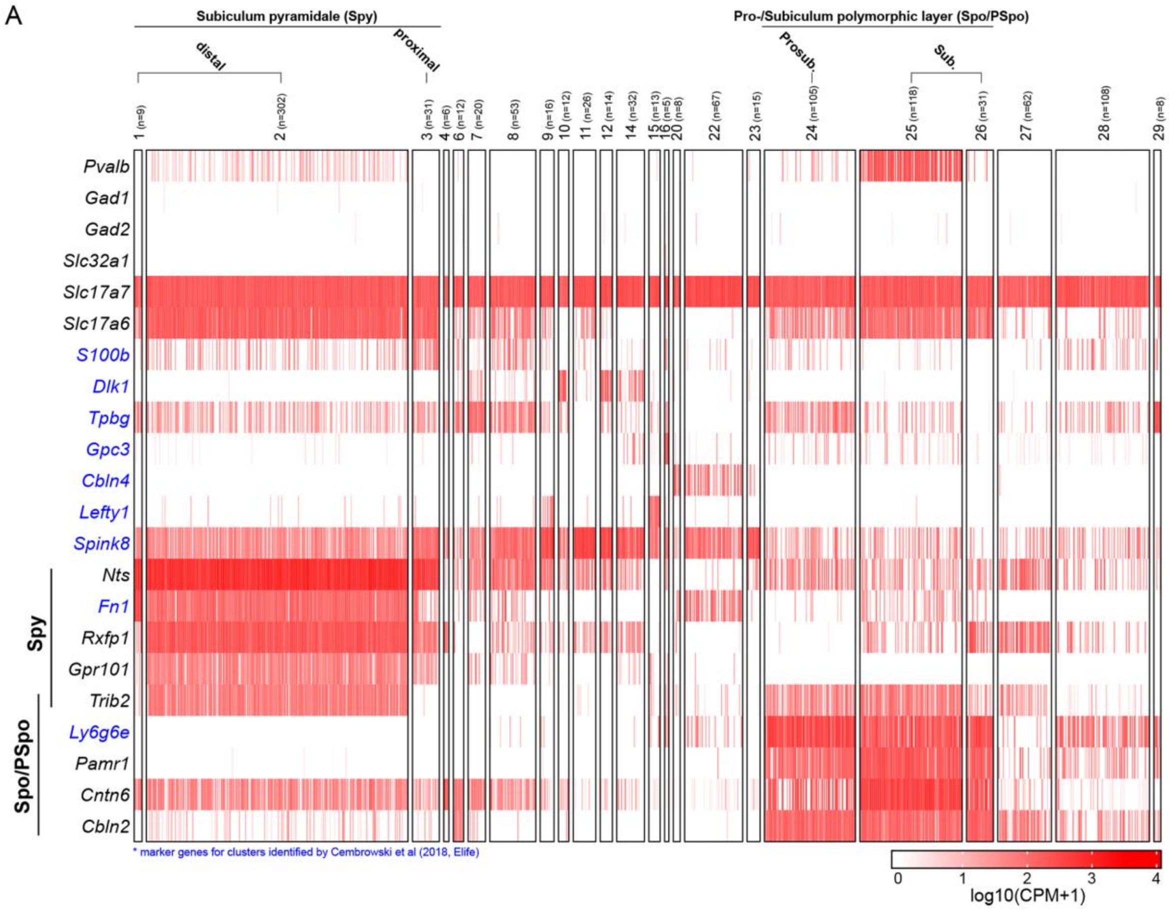


**Figure S11. PV-expressing principal cells map onto two different transcriptomic clusters in the distal subiculum.** Selection of genes that are differentially expressed in the clusters 1-29 obtained from the hierarchical re-clustering (RC1-29) of cells from the subiculum and prosubiculum, performed by Ding et al. 2020. The marker genes identified by Cembroswki et al., 2018, are shown in blue.

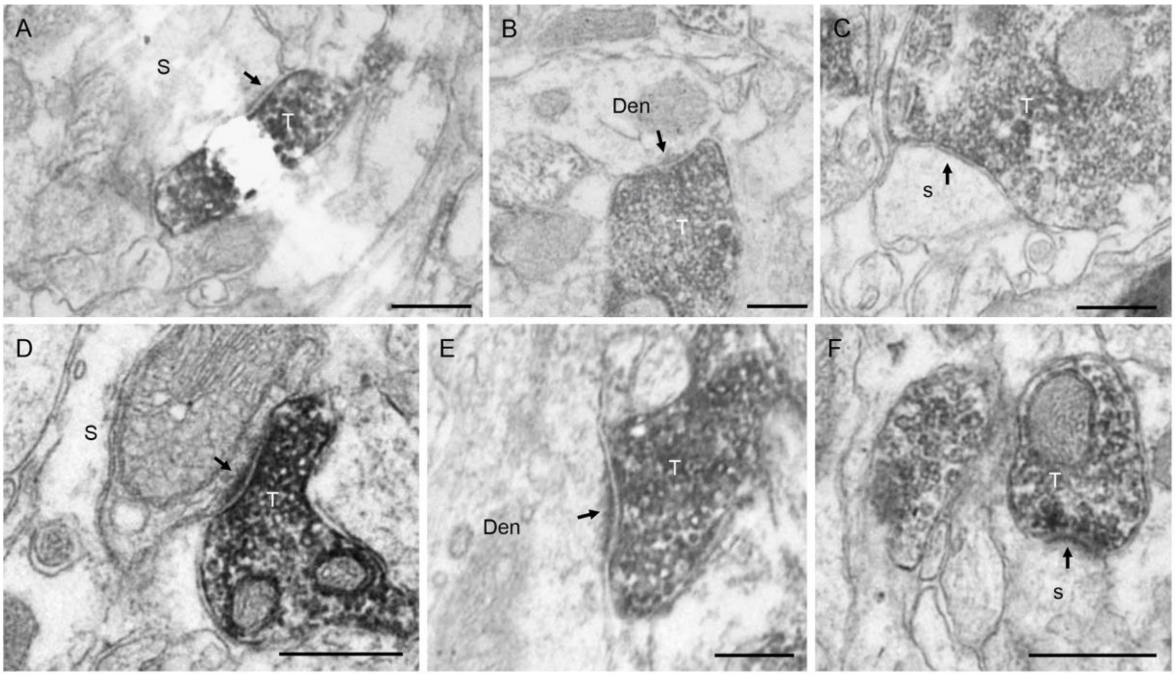


G

**Electronmicrographs**

| | | | | | | |
|---|---|---|---|---|---|---|
| **Synapses** | Number | | Location | | | Mice |
| | | | Soma | Shaft | Spine | |
| | 25 | ratio (number) | 0.08 (2) | 0.68 (17) | 0.24 (6) | 4 |
| **Colocalization** | Number GFP+ boutons | Colocalization ratio | Location | | | Mice |
| | | | Soma | Shaft | Spine | |
| VGLUT2+ | 46 | 0,5 | – | 0,81 | 0,19 | 3 |
| **Size GFP+ VGLUT2+ boutons** | Boutons number | Cross section areas of theG FP+ boutons (μm²) | | | | Mice |
| | | Mean | s.e.m. | Min | Max | |
| | 16 | 0,92 | 0,17 | 0,11 | 2 | 3 |
| **Size of contacted dendrites** | Number of dendrites | Diameter size (μm) | | | | Mice |
| | | Mean | s.e.m. | Min | Max | |
| | 14 | 0,84 | 0,24 | 0,29 | 2,69 | 3 |

**Neurobiotin-filled thalamic neurons**

| | | | | | | |
|---|---|---|---|---|---|---|
| **Dendrites size** | Proximal dendrites diameter (μm) | | Distal dendrites diameter (μm) | | Cells | Paired t-student |
| | Mean | s.e.m. | Mean | s.e.m. | | |
| | 0,96 | 0,08 | 0,64 | 0,08 | 3 | P=0.013 |

**Figure S12. Quantitative analysis of GFP+ terminals from the subiculum in the anteroventral thalamus.** A-F) GFP-positive (peroxidase reaction end-product) axon terminals (T) making symmetric (A-C) and asymmetric (D-F) synapses (arrows) on somata (S in A, D), dendritic shafts (Den in B, E) and dendritic spines (s in C, F) of postsynaptic neurons in the AV nucleus of the thalamus. Scale bars, 200 nm. G) Table reporting the distribution and size of GFP+ terminals. The upper part of the table reports data from electron microscopy. Cross section area of boutons and size of contacted dendrites were measured on randomly chosen samples. The lower part reports measurements of dendritic size performed on confocal images of reconstructed cells (see Fig. 4).

| | AV responding cells | | | | AV non-responding cells | | | |
|---|---|---|---|---|---|---|---|---|
| **Subthreshold** | **Mean** | | **s.e.m.** | **n** | **Mean** | | **s.e.m.** | **n** |
| Resting membrane potential (mV) | -69.82 | ± | 1.58 | 13 | -66.35 | ± | 2 | 11 |
| $R_N$ (MΩ) | 544.4 | ± | 30.65 | 14 | 434.9 | ± | 31.37 | 13 |
| I Holding (pA) | 14.83 | ± | 3.84 | 14 | 2 | ± | 6.65 | 13 |
| **Suprathresold, at rheobase** | | | | | | | | |
| Full-width at half-maximum (FWHM) (ms) | 1.26 | ± | 0.07 | 15 | 1.21 | ± | 0.06 | 6 |
| Frequency at double the rheobase (Hz) | 33.68 | ± | 4.64 | 8 | 47.86 | ± | 9.59 | 7 |
| Number of AP during rebound burst | 3.35 | ± | 0.4 | 14 | 3.42 | ± | 0.7 | 7 |

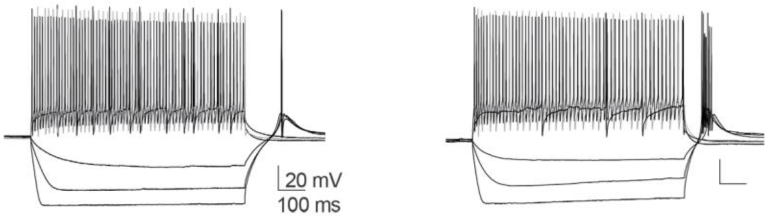


**Supplementary Table 1. Basic electrophysiological properties of neurons in the antero-ventral thalamus.** Values are shown separately for neurons that responded to light stimulation with EPSPs and those that did not. Voltage responses to increasing current injection of a responding (left) and a non-responding (right) neuron are shown below the table.
